# Unveiling the chemical and behavioural ecology of *Tribolium castaneum* (Herbst, 1797) in wheat flour: Alterations in flour metabolic content and the role of chemical cues in modulating beetles’ behaviour and regulating population growth

**DOI:** 10.1101/2024.10.01.616151

**Authors:** Subhadeep Das, Sourav Manna, Oishika Chatterjee, Riya Saha, Oishee Janet Sarkar

## Abstract

*Tribolium castaneum* (Herbst, 1797) (Coleoptera: Tenebrionidae) is a notorious secondary pest of stored grains and flours, accounting for 10–15% of global annual losses in stored products. This study first evaluated the impact of *T. castaneum* infestation on the nutritional parameters of stored wheat flour. Infested flour exhibited reductions in total soluble sugars, triacylglycerol levels, and unsaturated fatty acids, particularly Oleic and Linoleic acid, while secondary metabolites such as phenolics, flavonoids, and tannins were increased. Behavioural assays revealed that adult beetles were significantly more attracted towards infested flour odour due to the presence of 4,8-Dimethyldecanal (4,8-DMD), an aggregation pheromone of T. castaneum. Another key compound in infested flour, 1-Pentadecene, derived from the beetles’ cuticular wax, was found to influence adult beetle behaviour. At higher concentrations, 1-Pentadecene acted as a repellent for the species. Molecular docking studies suggested that 1-Pentadecene competes with 4,8-DMD for binding to the odorant receptor TcOR1, which could potentially inhibit aggregation behaviour and reduce population growth. The docking analysis revealed that 1-Pentadecene interacts with the same binding pocket on TcOR1, sharing key amino acids (MET49, PHE69, THR72, GLN73, THR134, VAL135, TYR138, LEU154, ILE177, GLY180, ALA181, TYR286) and forming similar intermolecular bonds, such as a pi-alkyl bond, with a binding affinity (ΔG = -5.7) comparable to 4,8-DMD (ΔG = -5.8). Additionally, beetles reared in wheat flour containing higher concentrations of 1-Pentadecene exhibited lower population growth compared to those reared in flour with lower concentrations of this compound.

## 1. Introduction

Wheat is a major staple food globally, with its production for the years 2023–24 estimated at approximately 785 million metric tons (Xu et al., 2024). In India alone, the wheat production is estimated to be 1129.25 lakh metric tons, which accounts for more than 30% of the total food grain production of this crop in India (DES, 2024). Despite such high production rates, both pre- and post-harvest losses pose significant threats to the productivity and supply of wheat products to consumers (Atta et al., 2020). It is estimated that around 15% post-harvest losses occur in the field, followed by 13-20% losses during processing and 15-25% losses during storage (Abass et al. 2014; Manandhar et al. 2018). The main post-harvest problem is insects, accounting for 5–15% of the loss at the storage stage (Negi et al., 2021).

The red flour beetle, *Tribolium castaneum* (Herbst, 1797) (Coleoptera: Tenebrionidae), is a cosmopolitan pest of stored grains, and is recognized as one of the major secondary insect pests (Đukić et al., 2020; Negi et al., 2021), which infests a wide variety of products, such as grains, flours, peas, beans, nuts, dried fruits and spices, and is responsible for 15–20% damage to the abovementioned commodities (Atta et al., 2020). *T. castaneum* degrades the nutritional value of stored products and often releases its own metabolites in the stored foods and thus reduces the quality of the stored products (Shamjana et al., 2021). Multiple lines of evidence suggesting that the metabolites secreted by *T. castaneum,* such as Benzoquinones (BQ), may pose significant health risks to consumers, with potential toxic, allergic, and carcinogenic effects in both humans and experimental animals (El-Desouky et al., 2018; El-Mofty et al., 1992; Lis et al., 2011; Pappas and Morrison, 1995).

However, before understanding the consequences of *T. castaneum* infestation, it is crucial to first explore how *T. castaneum* can identify its food sources and become inoculated. The inoculation and dispersal of *T. castaneum* are influenced by various external stimulus including visual cues (Semeao et al., 2011) and olfactory cues (Đukić et al., 2020). Few studies have suggested that adults of *T. castaneum* can respond to various visual stimuli, including colours and shapes. Semeao et al., (2011) suggested that *T. castaneum* prefers black objects over white ones when placed against a white background, and this preference intensifies as the height of black objects increases. In addition to visual cues, olfactory signals, including airborne volatile organic compounds (VOCs) and low-volatile compounds such as hydrocarbons, perceived by the chemosensory sensilla on the antennae, can also influence the localization behaviour of *T. castaneum*, as has been observed in many other insect species (Ai et al., 2022; Renou and Anton, 2020; Suh et al., 2014; Trebels et al., 2021). Additionally, the odours emitted by same food source, but with varying degrees of processing and quality, can differentially influence the attractiveness and food preferences of *T. castaneum*. The beetles exhibited a greater attraction to volatiles emitted by the wheat endosperm, germ, flour and bran than to the volatiles from whole wheat grains. Notably, wheat germ extracts were found to be more attractive for *T. castaneum* than extracts from other wheat fractions (Seifelnasr et al., 1982). On a similar note, the author also showed that, volatiles from normal and fermented millet flour were more attractive to *T. castaneum* than those from whole millet kernels (Seifelnasr et al., 1982). Furthermore, other studies reveal that odours from pre-infested stored products positively influence the attractiveness of *T. castaneum, in* compared to uninfested products. (Athanassiou et al., 2016; Stevenson et al., 2017).

Once infested by *T. castaneum,* the infested flour might have different odour profile. Some of these alterations are due to the breakdown of flour metabolites, which results in altered odours (Shamjana et al., 2021), while others are attributed to secretions from *T. castaneum* body, as mentioned earlier. Besides BQ, 4,8-Dimethyldecanal (4,8-DMD; aggregation pheromone of genus *Tribolium*) is another such compound, which accumulates in flour over time and has been reported to enhance the attractiveness of food for *T. castaneum* (Đukić et al., 2020; Suzuki, 1980). Studies showed that when synthetic 4,8-DMD was added to the food, the food was more appealing to the beetles than the scents of food or pheromone alone. (Campbell, 2012; Dissanayaka et al., 2020, 2018; Phillips, 1997). Apart from 4,8-DMD and BQ, there are other odorants such 1-Pentadecene, 1-Hexadecene etc. that are contributed to stored products from *T. castaneum* secretions, and the concentrations of *T. castaneum* odorants in flour are often considered as good indicators of population density of *T. castaneum* (Pointer et al., 2021). Given the exclusivity of these odorants in stored flour, some authors have suggested that these compounds may serve as volatile markers for these pests and can aid in establishing quick detection techniques at low infestation stages in stored items (Han et al., 2023; Villaverde et al., 2007). However, to effectively use such volatiles as biomarkers for early detection and population monitoring of red flour beetle infestation in wheat flour, it is crucial to identify such volatiles that could serve as chemomarkers for *T. castaneum* infested flour. This includes distinguishing between odorants *released by T. castaneum* and those generated through the metabolic alterations of the wheat flour. Additionally, it is important to understand how these odorants influence the behavioural responses of *T. castaneum* and affect their food preferences. In this article, we aim to address the aforementioned questions and try to explore how the interplay of multiple odorants released by *T. castaneum* in wheat flour influences the beetle’s preference for infested flour. Additionally, we examined how these odorants shape and regulate the population size of *T. castaneum* in wheat flour.

## 2. Materials and methods

### 2.1. Insect Rearing and Experimental Setups

Whole grain wheat flour was purchased from a local mill and sterilized in a hot air oven at 60°C for 30 minutes to eliminate any existing eggs, pupae, larvae of *T. castaneum*, or other unwanted insects. Fifteen grams of wheat flour was then measured and placed into 100 ml round plastic containers. In each container, ten 7-day-old adult *T. castaneum* (5 males and 5 females) were introduced. The cultures were maintained under a 12-hour light:12-hour dark photoperiod at 25°C ± 1°C and 65% ± 3% humidity for three months. After that *T. castaneum* infested flours were collected and used for further study. For comparative analysis, fresh wheat flour was used as control. The beetles were also collected and used in subsequent behavioural and chromatographic studies.

### 2.2. Preparation of Samples for Spectrophotometric Analysis

50 mg of fresh wheat flour (WF) and *T. castaneum* infested wheat flours (TWF: 3 months post inoculation) were weighed and mixed with 1 ml of 90% ethanol (v/v) and 1 ml of 90% methanol (v/v). The mixtures were kept overnight at 4 °C, followed by centrifugation at 8000 rpm for 10 minutes. The supernatants were carefully separated; the ethanolic extract was subsequently used to determine the total soluble carbohydrate content, while the methanolic extract was used for estimation of total phenolic, hydrolysable tannin, terpenoid, and flavonoid contents. Four biological replicates of both WF and TWF were used for all biochemical analyses.

### 2.3. Analysis of Primary Metabolites

#### 2.3.1. Estimation of Total Soluble Sugars

The total soluble sugars were quantified using the (Yemm and Willis, 1954) method. For the analysis, 10 µl of the ethanol extract from each sample was diluted in 240 µl of ethanol. Then, 250 µl of this diluted sample was mixed with 750 µl of Anthrone reagent (prepared by dissolving 200 mg of Anthrone in 100 ml of diluted H_2_SO_4_). The mixture was heated in boiling water for 10 minutes. After cooling to room temperature, the total soluble carbohydrate content was determined at 620 nm wavelength using UV-VIS spectrophotometer (UV-2600, Shimadzu, Japan). The absorbance was quantified using sucrose standards for preparing the standard curve.

#### 2.3.2. Estimation of Total Soluble Protein

200 mg of both WF and TWF samples were mixed in 1 ml of 0.1 M PBS (pH 7.0). The supernatant was collected after centrifugation three times at 8000 rpm in 4°C for 20 min to determine the protein content using the Bradford ’s method (Bradford, 1976). 200µl of sample extract was mixed with 1ml of Bradford reagent, kept for 15 mins and the absorbance was measured at 595 nm in the spectrophotometer. The concentrations of the samples were calculated by using BSA standard curve.

#### 2.3.3. Estimation of Total Lipid and Fatty acid Analysis

Bligh and Dyer’s method were used to extract lipids from both WF and TWF (Bligh and Dyer, 1959). For gravimetric estimation of lipids, an aliquot of chloroform extract was obtained in a pre-weighed glass vial. With the help of a N2 stream, the solvent was thoroughly evaporated, and the residue was once again precisely weighed and expressed in mg/g.

Fatty acid analysis was done using another aliquot of chloroform phase. Through acid catalyzed esterification (Christie, 1993; Poddar-Sarkar, 1996), fatty acid methyl esterification (FAME) was done to the fatty acids found in the infested and uninfested wheat. Following the extraction of FAME using n-Hexane, it was then dried using anhydrous sodium sulphate (Na_2_SO_4_). The n-Hexane was reduced in volume under a nitrogen flow and then analysed using Gas Chromatography-Mass Spectrometry (GC-MS).

### 2.4. Analysis of Secondary Metabolites

#### 2.4.1. Estimation of Total Phenolic content and Hydrolysable Tannins

The total phenolic content (TPC) was quantified via the methodology outlined by Ainsworth and Gillespie (Ainsworth and Gillespie, 2007). According to this procedure, 50 µL of the methanolic extract was first diluted with 50 ml of 90% methanol (v/v). The diluted methanolic extract was then combined with 200 µL of a 10% (v/v) Folin-Ciocalteu solution. The mixture was vigorously swirled and left in the dark for 5 minutes. Next, 800 µL of sodium carbonate (700 mM) was added to each tube, followed by incubation of the reaction mixture in the absence of light for 2 hours. Absorbances at 765 nm and 738 nm were measured using a UV-VIS spectrophotometer (UV-2600, Shimadzu, Japan) for TPC and hydrolysable tannin content measurements, respectively, with 1 ml of each reaction mixture. Standard curves were constructed for the determination of TPC and hydrolysable tannins using gallic acid equivalents (GAE) and tannic acid equivalents (TAE), respectively

#### 2.4.2. Estimation of Total Flavonoid content

The total flavonoid content was estimated by following the modified protocol of Csepregi et al. (2013). First, 300 µl ddH2O was added to the 100 µl of methanolic extract. Subsequently, 30 µl of 5% Sodium nitrite was mixed in and the mixture was allowed to stand at room temperature for 5 minutes. After that, 30µl of 10% aluminium chloride was added, mixed and kept at room temperature for 5 min. At last, 200 µl of 1 Mm NaOH was added, the mixture was shaken and diluted with 340 µl of ddH2O and allowed to stand for 15 minutes prior estimation. Total absorbance was measured at 510 nm, using UV-VIS spectrophotometer. Total flavonoid content was calculated based on the regression equation derived from catechin standards at 510 nm.

#### 2.4.3. Estimation of Total Terpenoid content

Estimation of total terpenoid was done by following protocol of Łukowski et al. (2022). At first, 1.5 ml Chloroform was added to the 200 µl methanolic extract of samples and was vortexed for 2 minutes. Then 100 µl of conc. H2SO4 was added to the reaction mixture. After 2 hours of incubation in the dark, 50µl of brown residue was taken and 950 µl methanol was added to the residue. UV-VIS spectrophotometer was used to measure the absorbance at 538 nm. The standard curve for total terpenoid content quantification was made using linalool at 538 nm.

### 2.5. Analysis of Volatile Organic Compounds (VOCs) of WF and TWF, T. castaneum body odours and cuticular hydrocarbons (CHCs)

To extract the VOCs from flour samples, 100 mg of WF and TWF were separately mixed with 1 mL of DCM. The mixture was thoroughly vortexed and left to settle for 24 hours. Once the DCM phase became visibly separated, 500 μL of the DCM extract were collected from both samples. The extracts were then concentrated to a final volume of 100 μL under a stream of nitrogen. For the analysis of the VOC profile, 1 μL of the DCM extract was injected into the GC-MS. For the methanolic extract, 100 mg each of WF and TWF were separately mixed with 1 mL of methanol. The mixtures were vortexed and left to settle for 24 hours.

After the methanol phase was separated, it was evaporated to dryness under a nitrogen stream. Subsequently, 100 μL of DCM was added to the dried methanolic extract, from that 1ul of sample was inject to further analysis. To validate the findings from the WF and TWF solvent extracts, another sampling technique, Headspace Solid Phase Micro Extraction (HS-SPME), a solvent-free method known for its high sensitivity in volatile sampling, was also employed. 100 mg of WF and TWF were placed in 20 ml crimp headspace vials (Agilent), and the vials were sealed airtight using a manual vial crimper (Agilent). HSVs were adsorbed for one hour by attaching a 1 cm 50/30 µm divinylbenzene/carboxen/polydimethylsiloxane [(DVB/CAR/PDMS); (Supelco, USA) stableflexTM, 24 Ga] SPME fiber with the help of a manual assembly holder (Supelco, USA) over the glass vials. The adsorbed volatiles were then desorbed by inserting the fiber into the GC-MS injector port for further analysis.

To extract *T. castaneum* body odour, 10 adult beetles were gently crushed in 1 mL of DCM. Then the DCM extracts were concentrated to 50 µl with the help of N2 stream, and then 1 µl was injected into GC-MS for analysis.

To extract cuticular hydrocarbons (CHCs), 10 adult beetles were dipped in n-Hexane (1 mL) for one minute. The cuticular hydrocarbons in n-Hexane were concentrated to 50 µl under a mild nitrogen stream, and then 1 µl was injected into GC-MS for analysis.

### 2.6. GC-MS parameters for chemical analysis

Thermo Fisher Trace 1300 (Thermo Scientific, Milan, Italy) GC-MS was performed with a triple-quad mass spectrometer (Thermo MS-TSQ 9000) equipped with a wall-coated open tubular column (WCOT), TG-5 MS (30 m × 0.25 mm × 0.25 μm) that was linked to a 10 m Dura guard capillary column. GC-MS analysis was performed with the front inlet temperature set to 250°C. Helium, with a purity of greater than 99.99%, served as the carrier gas at a flow rate of 1 mL/min.

#### 2.6.1. GC-programme for analysis of FAME

The initial oven temperature was set at 70°C with 1 min hold time, followed by a ramp increase at a rate of 5°C/min until reaching 260°C. A final hold at 260°C was maintained for 10 minutes. Total run time was 49 minutes.

#### 2.6.1. GC-programme for analysis of DCM and Methanolic extract and HSVs of WF and TWF

The initial oven temperature was set at 50°C with 1 min hold time, followed by a ramp increase at a rate of 7.5°C/min until reaching 200°C. A final hold at 200°C was maintained for 4 minutes. Total run time was 25 minutes.

#### 2.6.2. GC-programme for cuticular wax

The initial oven temperature was set at 50°C with 1 min hold, followed by a ramp increase at a rate of 6°C/min until reaching 300°C with a final hold of 8 minutes at the end. Total run time was 51 minutes

#### 2.6.3. GC programme for body extracts

The initial oven temperature was set at 50°C with 1 min hold time, followed by a ramp increase at a rate of 7.5°C/min until reaching 280°C. A final hold at 280°C was maintained for 3 minutes. Total run time was 35 minutes.

For all of the above programme, the transfer line temperature of the mass spectrometer was maintained at a constant 250°C, and the ion source temperature was set at 220°C. The mass spectrometer operated in electron ionization (EI) mode with an electron energy of 70 eV.

The data were processed using “Chromeleon Chromatography Data System (CDS) software” from Thermo Fisher Scientific, and the individual VOCs identified by matching m/z spectra with NIST 2020 mass spectral library, and Linear retention indices (LRIs) with the available unbranched alkanes (C8-C40) standards mixture (Sigma, Lot no: 40147-U). The LRI for any specific compound was estimated as:

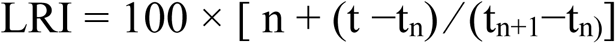

[Where n corresponds to the carbon number of the n-alkane that immediately precedes the target compound. t is the retention time of the compound, whereas t_n_ and t_n+1_, represent the retention times of the n-alkanes that directly precede and follow the VOC, respectively. These retention times are determined under identical experimental conditions, using the same temperature gradient and chromatographic column setup.].

The identification of FAME was done by comparing with authentic combination of 37 FAME and PUFA (Supelco, Lot No: LB80556 and LB77207, USA).

For analysis of GC-MS data, relative abundance of each compound was calculated by normalising its Total Ion Count (TIC) area with the total TIC area of all detected compounds.

### 2.7. Bioassays

#### 2.7.1. Bioassay 1: Effect of WF and TWF extract on T. castaneum in experimental arena

To assess the influence of various extracts from WF and TWF, a choice arena bioassay was conducted. Odorants from 100 mg of WF and TWF were extracted using 1 ml of dichloromethane (DCM). From each extract, 20 µl was applied onto a fresh Whatman filter paper disc (1 cm diameter), along with a blank DCM control. The treated discs were placed in a triangular arrangement inside a glass petri dish (9 cm diameter). Ten adult beetles (5 males and 5 females) were individually released at the centre of the petri dish, and their preference for the discs was observed. Beetle’s preferences for each disc were recorded. The experiment was repeated 10 times using different sets of beetles. Additionally, after each beetle was released, its position within the arena was monitored at 30-second intervals for 5 minutes to determine which area of the petri dish the beetle preferred to spend more time. A similar bioassay was conducted using methanolic extracts of WF and TWF (Supplementary Fig. 1A).

These experiments aimed to determine whether TWF extracts were more attractive to *T. castaneum* and to identify which solvent extract elicited a stronger behavioural response in the beetles.

#### 2.7.2. Bioassay 2: Effect of WF and TWF extract on T. castaneum in Y-tube olfactometer

A Y-tube olfactometer assay was conducted to validate the results obtained from the arena bioassay. In this assay, 20 µl of dichloromethane (DCM) extracts from both TWF and WF samples were tested individually against a DCM blank. Additionally, another choice test was performed to compare beetle preference between DCM extracts of TWF and WF samples (Supplementary Fig. 1B).

For each trial, 10 adult beetles (with a 1:1 male-to-female ratio) were released individually at the open end of the Y-tube, allowing them 1 minute to choose a direction. Each experiment was repeated five times with different sets of beetles.

#### 2.7.3. Bioassay 3: Food preference of T. castaneum towards WF and TWF

To determine which type of flour *T. castaneum* prefers for colonization, a bioassay was conducted. A 9 cm diameter petri dish was divided into two equal halves using a 1 cm thick plexiglass barrier. One half of the dish contained 3 g of fresh wheat flour, while the other half contained 3 g of wheat flour that had been infested for three months. The middle 1 cm thick plexiglass platform was used as release site for the beetles, and 20 adult beetles (with a 1:1 male-to-female ratio) were released onto the platform. The setup was left undisturbed for 24 hours to allow the beetles to choose their preferred food substrate for colonization (Supplementary Fig. 1C). After 24 hours, the number of beetles present on each side of the dish was counted and recorded. The experiment was repeated five times.

#### 2.7.4. Bioassay 4: Effect of T. castaneum cuticular wax in determining beetle behaviour

To determine whether the addition of *T. castaneum* cuticular wax to the DCM extract of fresh wheat flour (WF) could elicit a response similar to that of the DCM extract of TWF, cuticular wax from *T. castaneum* was extracted using the previously described method. A 10 µl aliquot of cuticular wax extract was combined with 20 µl of WF DCM extract and applied in 1cm Wattman paper. The bioassay was conducted as outlined in Bioassay 1, with three distinct odour sources: odours from WF, odours from the combination of WF and *T. castaneum* cuticular wax, and a DCM blank (Supplementary Fig. 2A).

#### 2.7.5. Bioassay 5: Influence of varying concentrations of 1-Pentadecene in modulating beetle behaviour

A bioassay was conducted to investigate whether varying concentrations of 1-Pentadecene elicit differential responses in *T. castaneum*. Infested wheat flour (TWF) samples were analyzed for the presence of 1-Pentadecene and quantified using GC-MS, following the previously described method, with octanal as the internal standard. Three TWF samples were selected for the study where 1-Pentadecene (10 ng/g, 20 ng/g, and 30 ng/g) concentrations vary considerably, while other components, including 4,8-DMD, remained relatively constant (Supplementary Table 1). Odorants from 100 mg of these TWF samples were extracted in 1 ml of DCM, as described earlier. The behavioural response of adult *T. castaneum* of both sexes to these extracts was assessed using a Y-tube olfactometer assay (Supplementary Fig. 2B). For the bioassay, 20 µl of each extract from the selected TWF samples (with varying 1-Pentadecene concentrations) was tested against 20 µl of a DCM blank. 20 µl of DCM extract from WF was regarded as a control.

#### 2.7.6. Bioassay 6: Role of 1-Pentadecene in determining reproductive success of T. castaneum

The assay was conducted to investigate whether the presence of 1-Pentadecene affects the reproductive success of *T. castaneum*. Three separate setups were prepared, each consisting of a 10 ml plastic container holding 15 g of fresh wheat flour. To each container, TWF DCM extracts from the previous bioassay were added, containing varying concentrations of 1-Pentadecene (10 ng/g, 20 ng/g, and 30 ng/g) while maintaining nearly identical 4,8-DMD levels. The amount of TWF extract added was adjusted to achieve the desired final concentration of 1-Pentadecene in each setup, i.e., 10 ng/g, 20 ng/g, and 30 ng/g.

In each experimental setup, 10 adult beetles (with a 1:1 male-to-female ratio) were released into the container. The containers were kept undisturbed for 3 months under the same rearing conditions as mentioned earlier. After 3 months, the number of adult beetles in each container was counted and recorded for further analysis. The experiment was replicated for 3 times (Supplementary Fig. 3).

### 2.8. In Silico molecular docking analysis

TcOR1, the ortholog of the *Drosophila melanogaster* Or83b (DmOr83b) odorant receptor protein was chosen for docking study as macromolecule. The FASTA sequence (NCBI accession number-XP_015832943.2) was noted from NCBI and uploaded to the SWISS-MODEL website (protein structure homology-modelling server, accessible via the Expasy web server) to search for a template. First hit (alpha fold) was taken into consideration for homology modelling. OR protein was then downloaded as Protein Data Bank (PDB) format. Receptor preparation was carried out by using AutoDock Suite 4.2.6. The following procedures were involved in the preparation of receptors in AutoDock: the deletion of water molecules, the incorporation of polar hydrogens, and the addition of Kollman charges. Following the completion of the processing, the protein molecule was then exported from AutoDock in the format of Protein Data Bank, Partial Charge (Q), and Atom Type (T) (PDBQT). 3D structural data files (SDF) of volatile ligands were downloaded from “PubChem (https://pubchem.ncbi.nlm.nih.gov/)”, which were later converted into PDB and subsequently to PDBQT format using OpenBabelGUI (2.4.1) and AutoDock (4.2.6), respectively. Using PyRx 0.8 with the AutoDock Vina Plugin, docking was done between the volatiles (ligands) and ORs (protein molecules). 9 distinct ligand molecule postures were achieved by performing dockings while maintaining the exhaustiveness parameter at 8. Following docking, the ligand posture provided the lowest binding energy, and Discovery Studio Visualizer (v21.1.0.20298) was then used to visualise the interaction between the ligand and protein. The binding energy (ΔG) and the value of RMSD were used to determine which ligand posture was going to be the most effective in binding to the receptor.

### 2.8. Statistical Analysis and Graphical Representation

OriginPro 2021 (9.8.0.200) and Past4.10 have been used to perform statistical analyses and preparing graphical representations.

## 3. Results

### 3.1. Effects of T. castaneum infestation on nutritional parameters of wheat flour

The effect of *T. castaneum* infestation on wheat flour involves changes in the primary and secondary metabolites. There was a significant reduction in total soluble sugar content (p =0.03) in TWF compared to WF. Additionally, while a reduction in total lipid content was observed in TWF, this decrease was not statistically significant (p=0.053). In contrast, TWF exhibited a significantly higher total soluble protein content (p=0.03) compared to WF (Fig. 1 A-C). The Thin layer chromatography (TLC) analysis revealed a marked difference in the lipid classes between WF and TWF. In fresh wheat flour, the triacylglycerol (TAG) content was notably high, while in infested wheat flour, TAG levels were significantly reduced. Conversely, the free fatty acid (FFA) content was substantially higher in TWF compared to WF, where FFA levels were relatively low (Supplementary Fig. 4).

**Fig. 1.**
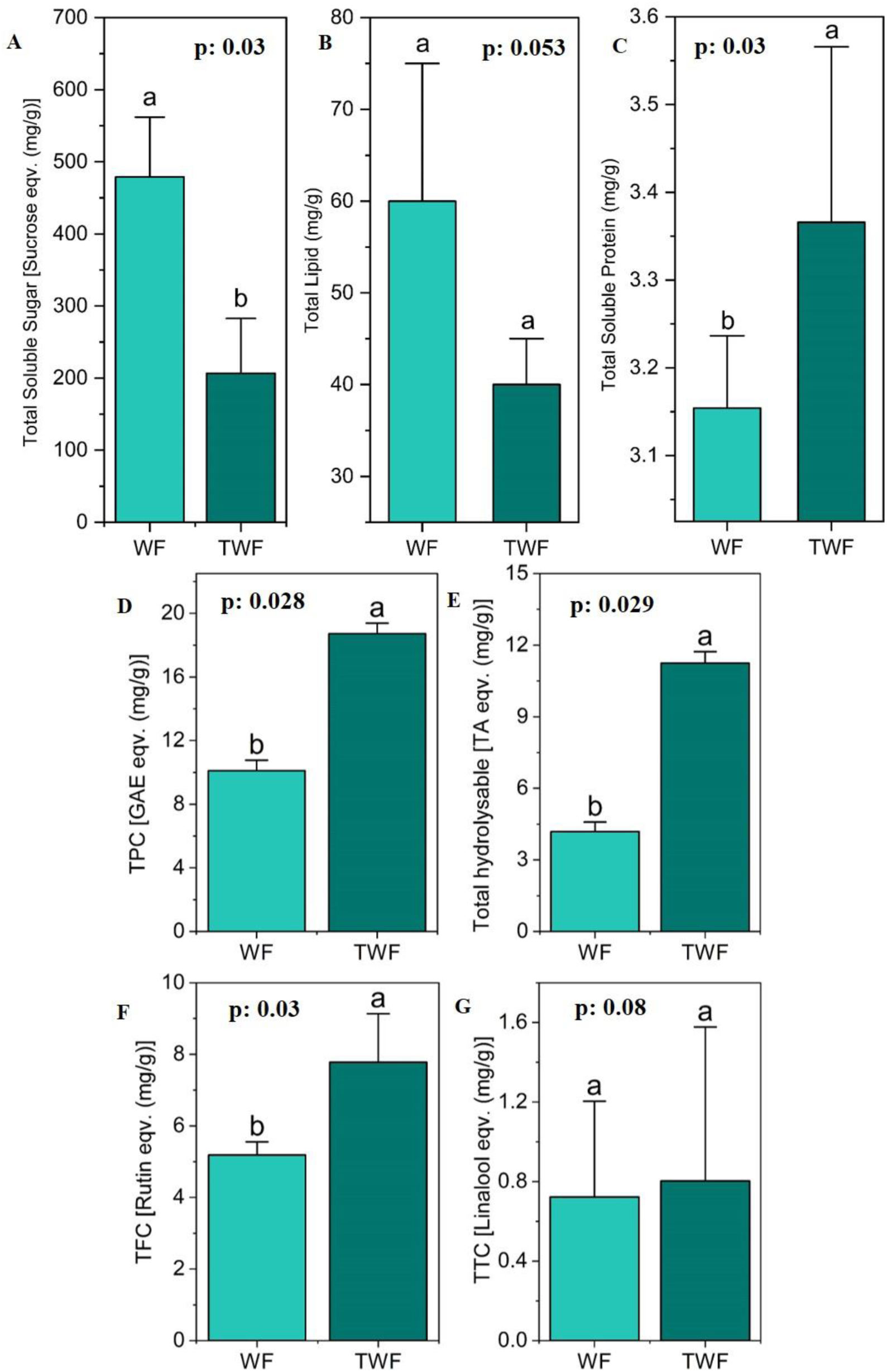
Bar diagram representing the changes in metabolite contents in WF and TWF: total soluble sugar (A), total lipid (B), total soluble protein (C), total phenolics (D), total hydrolysable tannins (E), total flavonoids (F), and total terpenoids (G). Statistical significance is denoted by p-values in the bar graph. Statistical significance is denoted by p-values, and results of the Mann-Whitney (U) test are indicated with letters above the bars. Bars that share same letter do not show any significant difference in metabolite content between WF and TWF.

Heatmap showing differences in fatty acid composition between WF and TWF (Fig. 2). The amount (relative %) of saturated fatty acids in TWF is higher than that of WF [WF_SFA_ = 41.07 ± 0.95%; TWF_SFA_ = 54.43 ± 0.15%], which is predominantly due to the amount of palmitic acid (C16:0) present in higher quantity in TWF than WF [TWF_C16:0_ = 49.73 ± 0.19%; WF_C16:0_ = 36.82 ± 2.14%], whereas the amount of unsaturated fatty acids (UFA) decreased in wheat flour after infestation [WF_UFA =_ 57.25 ± 3.04 %; TWF_UFA_ = 45.56 ± 0.15 %]. Linoleic acid (ω-6 fatty acid; C18:2n-6), one of the two essential dietary fatty acids, showed a decrease in amount following infestation [WF_C18:2n-6_ =36.49 ± 3.48 %; TWF_C18:2n-6_ = 23.42 ± 0.40 %] (Supplementary Table 2).

**Fig. 2.**
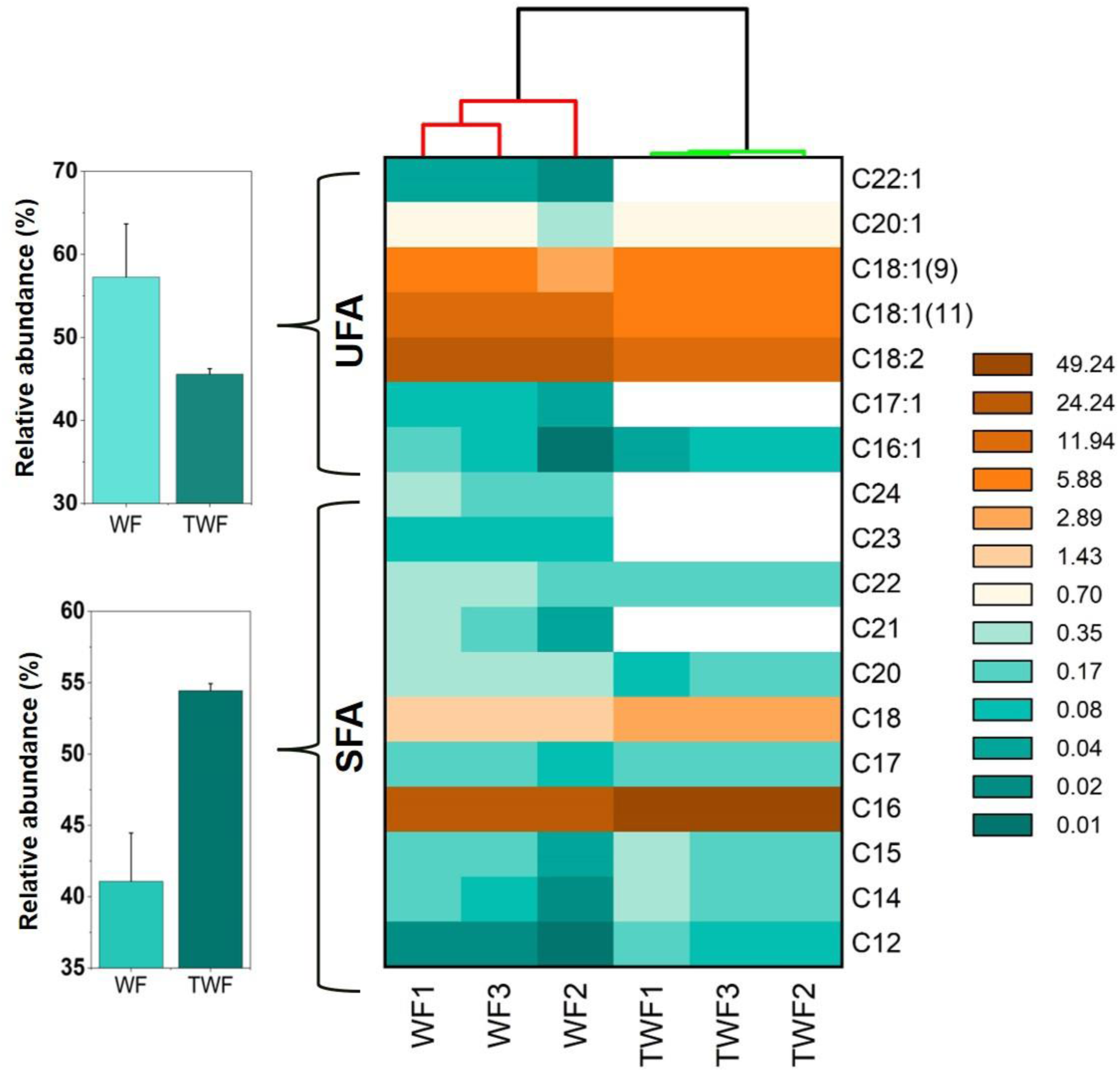
Heatmap illustrates the fatty acid profile analysis of WF and TWF. The color intensity represents the relative concentration of each fatty acid (in log scale). Clustering of samples depending on fatty acid profiles, represented by the dendrogram on top, shows the relationship between replicates within and between the WF and TWF samples.

Secondary metabolite contents such as phenolics, flavonoids, tannins, and terpenoids increased in TWF. TPC (p=0.028), TFC (p=0.029), and THT (p=0.03) content increased significantly in TWF compared to fresh WF. There was no significant increase in TTC content in TWF (Fig. 1 D-G).

### 3.2. Behavioural modulation activity of WF and TWF extract on T. castaneum

To investigate the role of olfactory cues in the food localization behaviour of *T. castaneum*, a choice bioassay was conducted using both DCM and methanolic extracts of WF and TWF (Supplementary Fig. 1A). The results indicated that the DCM extract elicited a stronger behavioural response compared to the methanolic extract. Both extracts from TWF acted as attractants for *T. castaneum* (Mean TWF_DCM_ = 7±2, Mean TWF_MeOH_ = 4±1), triggering similar behavioural responses. However, statistically significant attraction was observed only with the DCM extract (H= 24.07, p=1.201E-05) (Fig. 3A). In a spatial localization assay within the experimental arena (Supplementary Fig. 1A), adult beetles spent more time near the odour source for both the DCM and methanolic extracts. Notably, in the case of the DCM extract, adult beetles spent more time near the TWF odour source, whereas with the methanolic extract, they alternated more frequently between the WF and TWF odour sources (Fig. 3B).

**Fig. 3.**
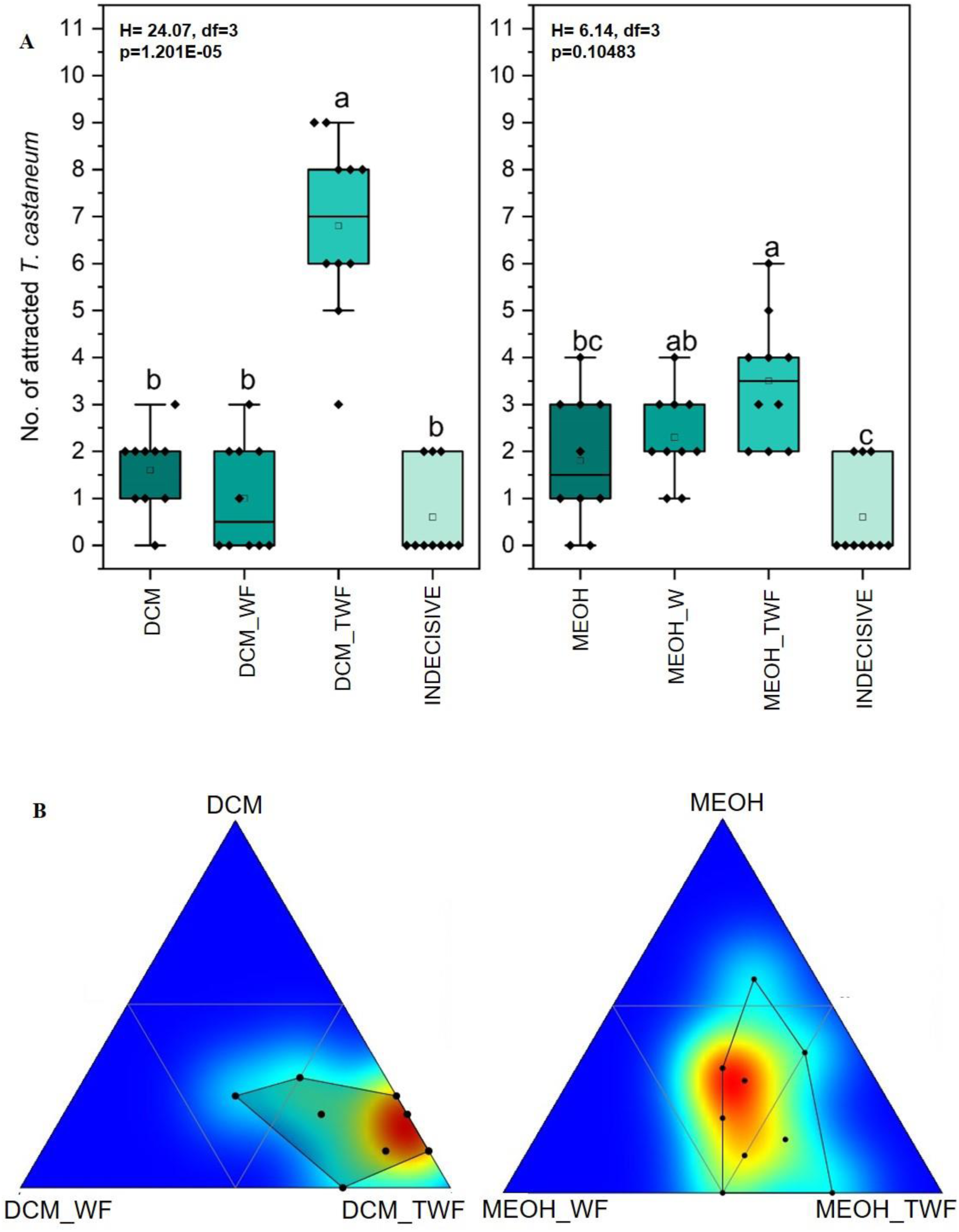
Box plot showing number of attracted beetles towards DCM (A Left) and MeOH extracts (A Right) of WF and TWF in choice bioassay. Significant changes (p<0.05) in beetle attraction were calculated using the Kruskal-Wallis test, followed by a Dunn’s post-hoc multiple comparison test. Letters above the box plots indicate statistical difference, where plots sharing the same letter show no significant difference in beetle attraction.

To further explore the role of odorants in the navigation of *T. castaneum* towards flour, Y-tube choice assays were conducted to assess the influence of WF and TWF odours on the odour preference of *T. castaneum* (Supplementary Fig. 1B). While testing response of adult beetles to WF and TWF DCM extracts individually, compared to a DCM blank, no statistically significant preference for WF odour was observed (U=3; z=1.365; p=0.172) (Fig. 4). In contrast, a significant preference for TWF odour was detected (U=0.5; z=2.032 p=0.042) (Fig. 4). In a separate Y-tube assay comparing WF and TWF odours, *T. castaneum* showed a significantly stronger attraction to TWF odour (TWF_DCM = 19 ± 2) compared to WF odour (WF_DCM = 11 ± 2) (U = 0, z = 2.165, p = 0.03) (Fig. 4).

**Fig. 4.**
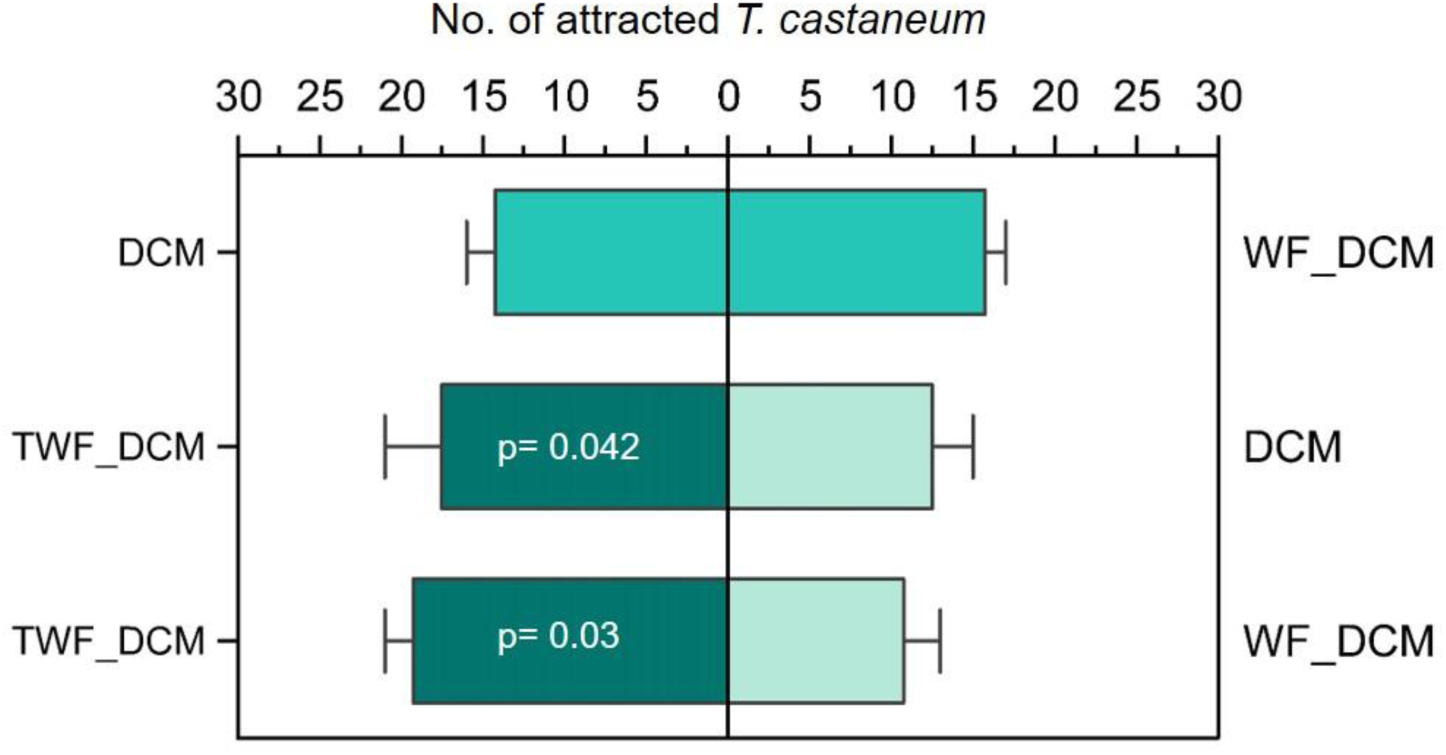
Two-choice olfactometer assay showing the number of *T. castaneum* individuals attracted to different treatments (DCM blank, WF_DCM and TWF_DCM). Statistical significance (p<0.05) obtained from Mann Whitney test is denoted by p-values in the bar graph.

### 3.3. Preference of T. castaneum towards WF and TWF

Further analysis was conducted to determine whether *T. castaneum* could differentiate between raw, uninfested flour and *T. castaneum*-infested flour (Supplementary Fig. 1C). In the first set of experiments, the adult beetles consistently preferred raw, uninfested flour (12±6) over infested flour (7±5) in all trials (U=7; z=1.057; p= 0.29) (Fig. 5). In the second experiment, where the preference between raw flour and raw flour supplemented with TWF DCM extract was tested, they showed a significant preference for the flour supplemented with exogenous odours (12±3) compared to the untreated raw flour (7±3); (U=2; z=2.121; p= 0.033) (Fig. 5).

**Fig. 5.**
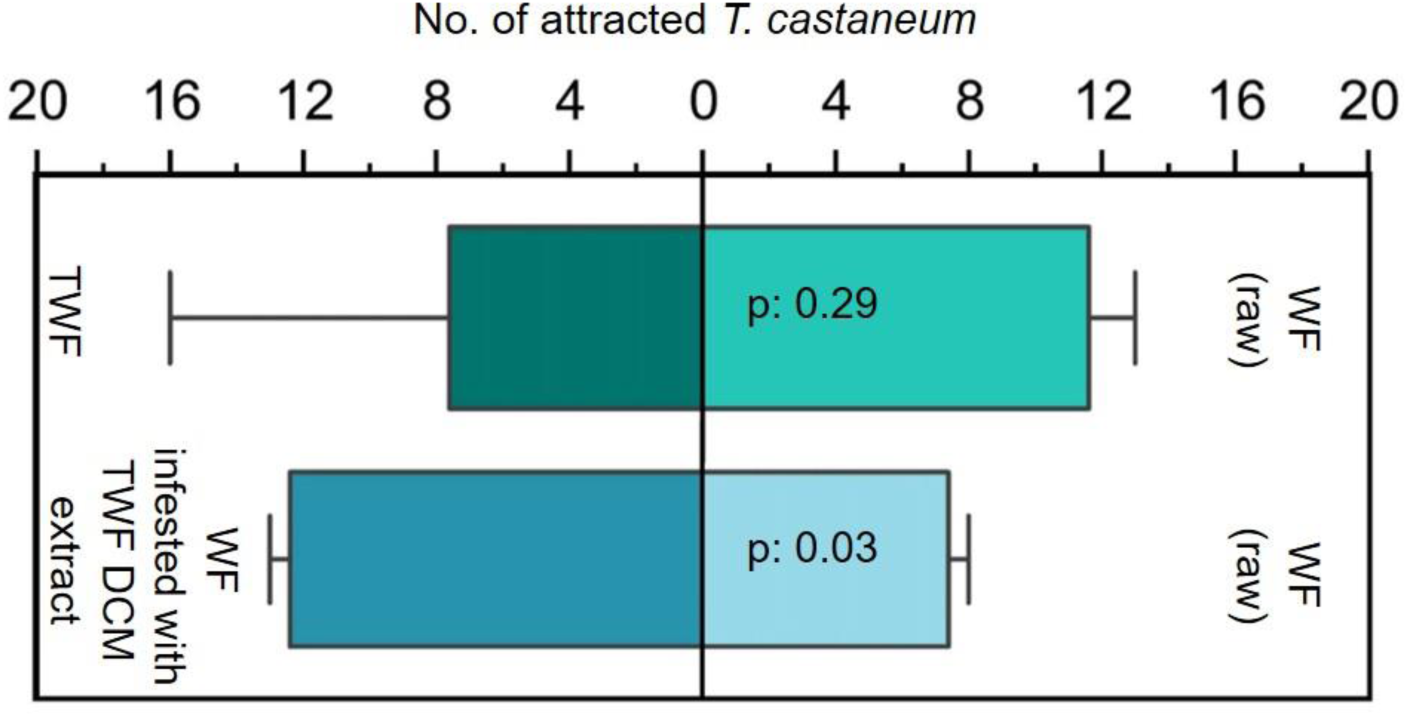
Preference of *T. castaneum* between fresh WF and fresh WF with DCM extracts of TWF. Statistical significance ((p<0.05) is calculated using Mann Whitney (U) test and denoted by p-values in the bar graph.

### 3.4 Characterization of odorant molecules present in WF and TWF

WF and TWF DCM extracts were found to contain 36 volatile organic compounds (VOCs) with various functional groups (Table 1, Fig. 6). Hydrocarbons represented the most abundant group of VOCs in both WF and TWF, comprising 52.1 ± 5.559% in WF and 53.46 ± 0.301% in TWF. In WF, the most abundant VOC was Cyclopentene, 1,2,3,4,5-pentamethyl (11.3 ± 2.551%), followed by 1-Butanol, 3-methoxy (6.32 ± 1.167%). In contrast, the most abundant VOC in TWF was 1-Pentadecene (12.09 ± 0.198%), followed by cis-9-Tetradecen-1-ol (8.40 ± 0.577%). 26 VOCs were common to both WF and TWF, with no significant differences observed in their concentrations (Supplementary Fig. 5a). TWF contained ten distinct VOCs, including Acetophenone [0.20 ± 0.216%], Neral [1.43 ± 0.002%], 4,8-Dimethyldecanal (4,8-DMD) [1.74 ± 0.158%], 1-Pentadecene [12.09 ± 0.198%], 1-Heptadecene [2.04 ± 0.195%], 2-Heptanone [0.83 ± 0.17%], cis-9-Tetradecen-1-ol [8.40 ± 0.577%], 2,3,3-Trimethyl-1-hexene [0.59 ± 0.347%], Cyclohexanol, 1-[3-(1-pyrrolidinyl)-1-propynyl] [0.04 ± 0.051 %] and Nonadecane [1.62 ± 0.117 %] (Supplementary Fig. 6).

**Fig. 6.**
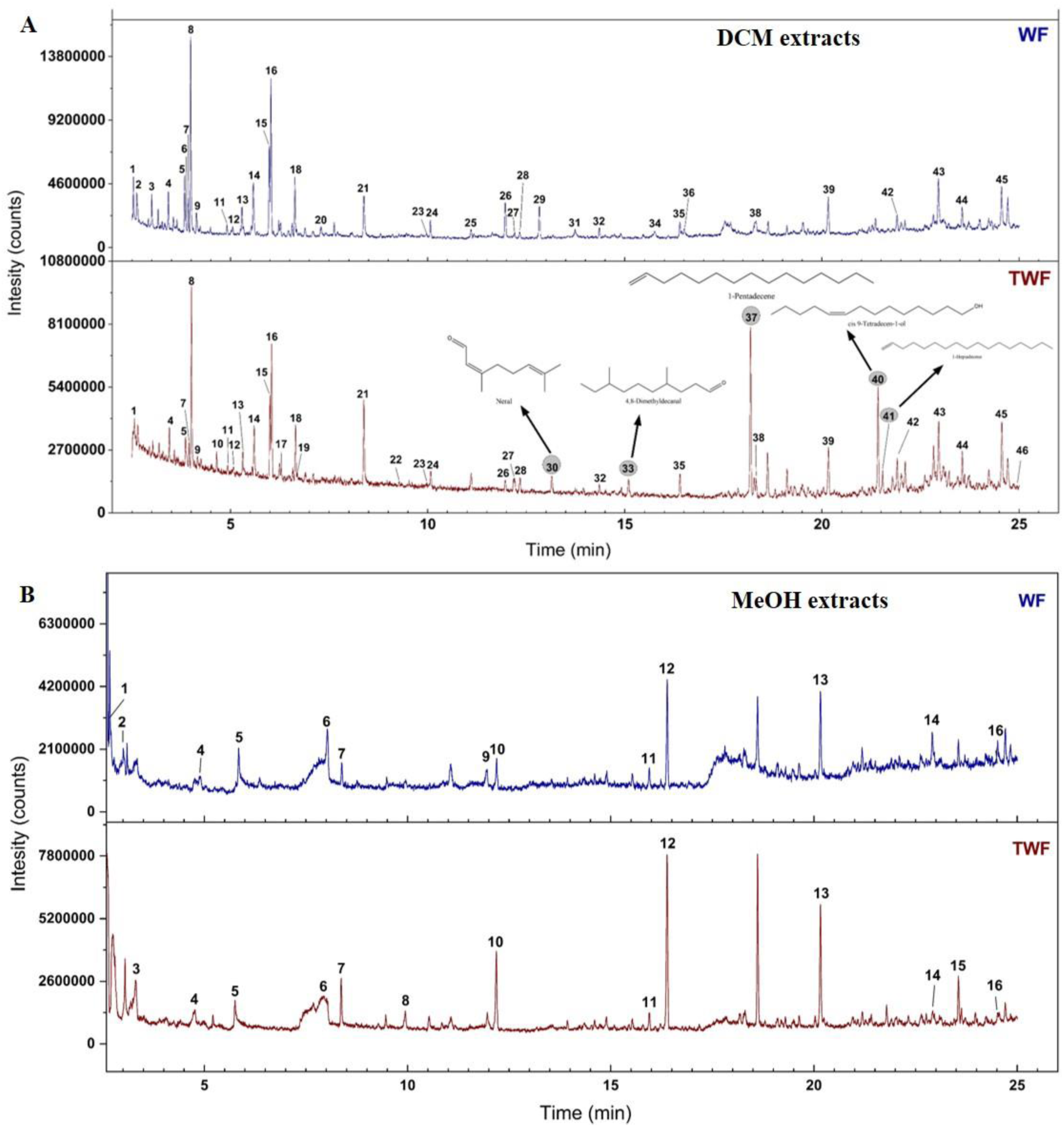
Gas chromatograms of DCM (A) and Methanolic (B) extracts of WF and TWF. **A.** 1. 3-Penten-2ol; 2. Heptane; 3. 3-Hydroxy-3-methyl-2-butanone; 4. 2-Hexanone; 5. Hexanal; 6. 3-Hexen-1-ol,(Z); 7. 1,1-Dimethyl-3-chloropropanol; 8. 2-Hexanol; 9. 1-Butanol,3-methoxy; 10. 2-Heptanone; 11. Ethylbenzene; 12. p-Xylene; 13. 2-sec-Butyl-3-methyl-1-pentene; 14. Cyclopentene,1,2,3,3,4-pentamethyl; 15. Cyclopentane,1,2,3,4,5-pentamethyl; 16. Cyclopentene,1,2,3,4,5-pentamethyl; 17. 2,3,3-Trimethyl-1-hexene; 18. m-Menthane; 19. Cyclohexanol,1-[3-(1-pyrrolidinyl)-1-propynyl]; 20. Hexanoic acid; 21. 1-Hexanol,2-ethyl; 22. Acetophenone; 23. Undecane; 24. Nonanal; 25. Acetic acid, 2-ethylhexyl ester; 26. Naphthalene; 27. Dodecane; 28. Decanal; 29. Carbonic acid, ethyl phenyl ester; 30. Neral; 31. Nonanoic acid; 32. Tridecane; 33. 4,8-Dimethyldecanal ; 34. Decanoic acid; 35. Tetradecane; 36. Vanillin; 37. 1-Pentadecene; 38. Pentadecane; 39. Hexadecane; 40. cis-9-Tetradecen-1-ol; 41. 1-Heptadecene; 42. Heptadecane; 43. Tetradecanoic acid; 44. Octadecane; 45. Pentadecanoic acid; 46. Nonadecane **B.** 1. Acetic acid; 2. 2-Propanone,1-hydroxy; 3. Propanoic acid, 2-methyl, ethyl ester; 4. 2-Pentanone,4-hydroxy-4-methyl; 5. Oxime, methoxy-phenyl; 6. Benzene,1,4-dichloro; 7. 1-Hexanol,2-ethyl; 8. Undecane; 9. Naphthalene; 10. Dodecane; 11. Propanoic acid,2-methyl,3 hydroxy-2,2,4-trimethylpentyl ester; 12. Tetradecane; 13. Hexadecane;14. Tetradecanoic acid; 15. Octadecane; 16. Pentadecanoic acid.

**Table 1.** List of identified VOCs from DCM and MeOH extracts of WF and TWF.

| Rt | Compounds | DCM extract |  | LRI<br>(TG5) |
| --- | --- | --- | --- | --- |
|  |  | WF (DCM<br>extract) | TWF (DCM<br>extract) |  |
| 2.576 | 3-Penten-2ol | ✓ | ✓ |  |
| 2.598 | Heptane | ✓ |  |  |
| 2.973 | 3-Hydroxy-3-methyl-2-butanone | ✓ |  | 728 |
| 3.137 | 2-Hexanone | ✓ | ✓ | 742 |
| 3.829 | Hexanal | ✓ | ✓ | 802 |
| 3.842 | 3-Hexen-1-ol,(Z) | ✓ |  | 803 |
| 3.897 | 1,1-Dimethyl-3-chloropropanol | ✓ |  | 806 |
| 3.973 | 2-Hexanol | ✓ | ✓ | 808 |
| 4.136 | 1-Butanol,3-methoxy | ✓ | ✓ | 820 |
| 4.607 | 2-Heptanone |  | ✓ | 847 |
| 4.897 | Ethylbenzene | ✓ | ✓ | 864 |
| 5.036 | p-Xylene | ✓ | ✓ | 871 |
| 5.274 | 2-sec-Butyl-3-methyl-1-pentene | ✓ | ✓ | 886 |
| 5.562 | Cyclopentene,1,2,3,3,4-pentamethyl | ✓ | ✓ | 902 |
| 5.965 | Cyclopentane,1,2,3,4,5-pentamethyl | ✓ | ✓ | 921 |
| 6.008 | Cyclopentene,1,2,3,4,5-pentamethyl | ✓ | ✓ | 923 |
| 6.247 | 2,3,3-Trimethyl-1-hexene |  | ✓ | 934 |
| 6.649 | m-Menthane | ✓ | ✓ | 951 |
| 6.659 | Cyclohexanol,1-[3-(1-pyrrolidiny)-1-propynyl] |  | ✓ | 953 |
| 7.33 | Hexanoic acid | ✓ |  | 985 |
| 8.356 | 1-Hexanol,2-ethyl | ✓ | ✓ | 1031 |
| 9.251 | Acetophenone |  | ✓ | 1070 |
| 9.956 | Undecane | ✓ | ✓ | 1103 |
| 10.068 | Nonanal | ✓ | ✓ | 1106 |
| 11.092 | Acetic acid, 2-ethylhexyl ester | ✓ |  | 1152 |
| 11.943 | Naphthalene | ✓ | ✓ | 1190 |
| 12.179 | Dodecane | ✓ | ✓ | 1202 |
| 12.333 | Decanal | ✓ | ✓ | 1229 |
| 12.793 | Carbonic acid, ethyl phenyl ester | ✓ |  | 1230 |
| 13.138 | Neral |  | ✓ | 1245 |
| 13.77 | Nonanoic acid | ✓ |  | 1274 |
| 14.355 | Tridecane | ✓ | ✓ | 1301 |
| 15.08 | 4,8-dimethyldecanal |  | ✓ | 1336 |
| 15.747 | Decanoic acid | ✓ |  | 1369 |
| 16.39 | Tetradecane | ✓ | ✓ | 1400 |
| 16.475 | Vanillin | ✓ |  | 1405 |
| 18.189 | 1-Pentadecene |  | ✓ | 1493 |
| 18.315 | Pentadecane | ✓ | ✓ | 1500 |
| 20.153 | Hexadecane | ✓ | ✓ | 1600 |
| 21.418 | cis-9-Tetradecen-1-ol |  | ✓ | 1671 |
| 21.535 | 1-Heptadecene |  | ✓ | 1673 |
| 21.899 | Heptadecane | ✓ | ✓ | 1700 |
| 22.965 | Tetradecanoic acid | ✓ | ✓ | 1763 |
| 23.549 | Octadecane | ✓ | ✓ | 1798 |
| 24.584 | Pentadecanoic acid | ✓ | ✓ | 1864 |
| 25.134 | Nonadecane |  | ✓ | 1901 |
| Methanolic extract |  |  |  |  |
| Rt | Compounds | WF (MeOH<br>extract) | TWF<br>(MeOH<br>extract) | LRI<br>(TG5) |
| 2.675 | Acetic acid | ✓ |  |  |
| 3.004 | 2-Propanone,1-hydroxy | ✓ |  |  |
| 3.315 | Propanoic acid, 2-methyl, ethyl ester |  | ✓ |  |
| 4.771 | 2-Pentanone,4-hydroxy-4-methyl | ✓ | ✓ |  |
| 5.757 | Oxime, methoxy-phenyl | ✓ | ✓ | 902 |
| 7.93 | Benzene,1,4-dichloro | ✓ | ✓ | 1016 |
| 8.369 | 1-Hexanol,2-ethyl | ✓ | ✓ | 1030 |
| 9.942 | Undecane |  | ✓ | 1101 |
| 11.958 | Naphthalene | ✓ |  | 1190 |
| 12.196 | Dodecane | ✓ | ✓ | 1201 |
| 15.949 | Propanoic acid,2-methyl,3-hydroxy-2,2,4-trimethylpentyl ester | ✓ | ✓ |  |
| 16.388 | Tetradecane | ✓ | ✓ | 1400 |
| 20.154 | Hexadecane | ✓ | ✓ | 1600 |
| 22.907 | Tetradecanoic acid | ✓ | ✓ | 1763 |
| 23.554 | Octadecane |  | ✓ | 1798 |
| 24.517 | Pentadecanoic acid | ✓ | ✓ | 1864 |

The table presents the retention times (Rt), compound names, and Linear Retention Indices (LRI) of the respective compounds. The LRI values are relative to C8-C40 n-alkanes and were obtained using a semi-standard nonpolar column (TG-5).

Compared to the DCM extracts, the methanolic extracts of WF and TWF contained fewer VOCs, with only 13 compounds identified (Table 1; Fig. 6). 9 VOCs were common to both WF and TWF, with no significant differences observed in their abundance (Supplementary Fig. 5b). No unsaturated hydrocarbons were detected in either flour. The most abundant compound in the methanolic extracts was Tetradecane, accounting for 12.83 ± 11.84% in WF and 23.60 ± 5.472% in TWF, followed by Hexadecane at 11.9 ± 10.988% in WF and 16.7 ± 1.364% in TWF. Headspace analysis revealed the presence of 19 and 26 headspace volatiles (HSVs) in WF and TWF, respectively, with various functional groups (Supplementary Table 3). In TWF, both 4,8-DMD and 1-Pentadecene were detected in the headspace, which were also present in the DCM extract and are known to play a pheromonal role in *T. castaneum*. Additionally, 6-Methyl-5-hepten-2-one was exclusively detected in the headspace of TWF and was not present in its DCM extract.

To identify the chemical markers associated with *T. castaneum*-infested wheat flour, principal component analysis (PCA) was performed. PC1 and PC2 accounted for 69.96% and 17.83% of the variance, respectively (Eigenvalues: PC1 = 95.16, PC2 = 24.25), distinguishing between WF and TWF samples based on the odorant molecules detected in the DCM extracts. Several volatiles were strongly correlated with TWF samples, with the highest loadings observed for 1-Pentadecene (0.672), cis-9-Tetradecen-1-ol (0.4688), 1-Heptadecene (0.1143), 2-Hexanol (0.1088), and 4,8-DMD (0.0964). In contrast, volatiles such as Hexanal (-0.1358), Carbonic acid ethyl phenyl ester (-0.1535), Cyclopentane, 1,2,3,4,5-pentamethyl (-0.154), Naphthalene (-0.1591), and Cyclopentene, 1,2,3,4,5-pentamethyl (-0.283) were more strongly associated with WF samples (Supplementary Table 4). Suggesting that there are a number of volatile compounds that make WF and TWF samples unique from each other.

### 3.5. Odorants of T. castaneum and its contribution in determining TWF odour

The analysis of the odour profiles of TWF, focusing on the impact of body odorants from T. castaneum, was conducted using GC-MS on both the beetles’ body extract and cuticular wax profiles. A total of 34 odorants were identified from the body extracts of TC, while 30 cuticular wax compounds were detected from the body surface (Table 2; Fig. 7A-B).

**Fig. 7.**
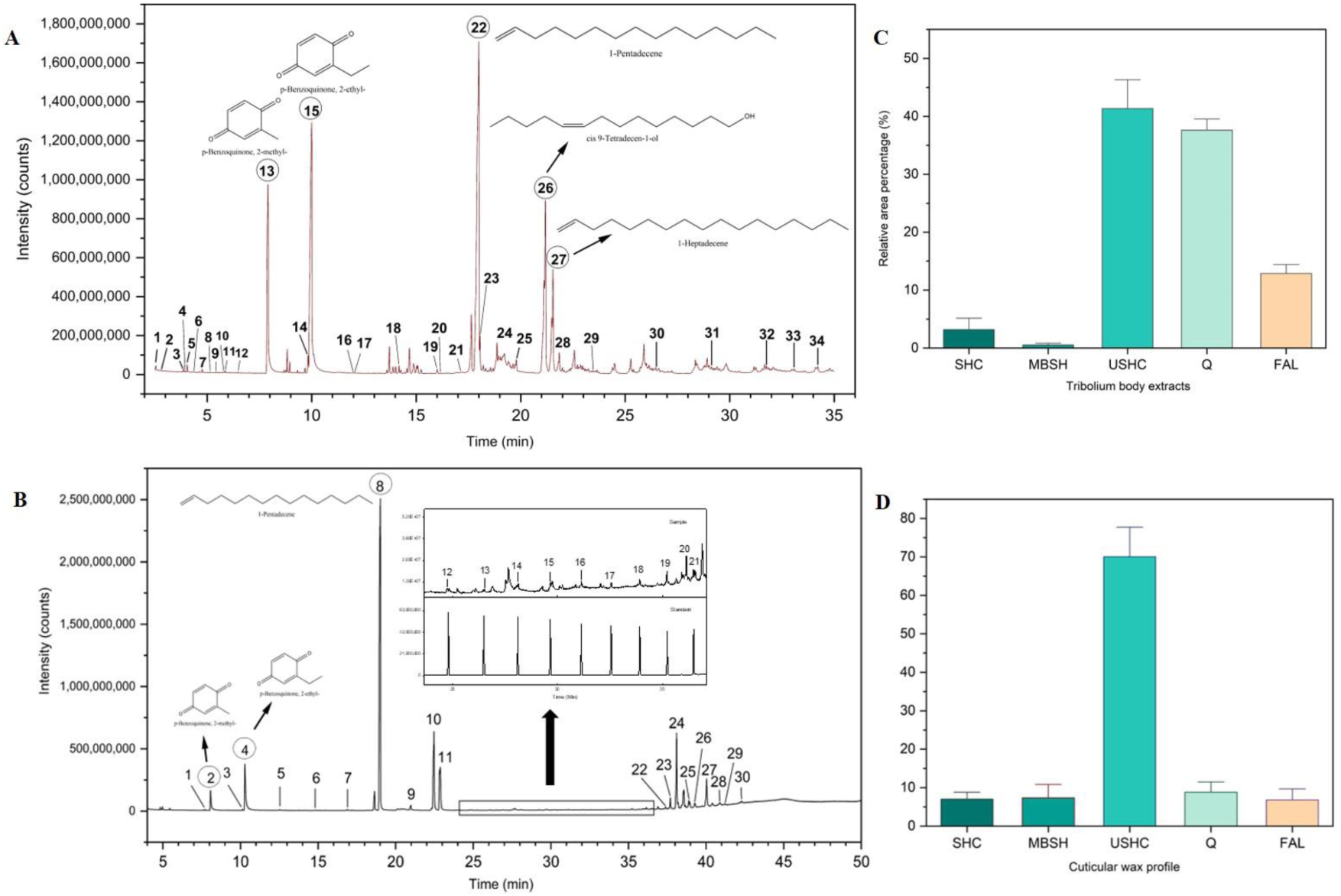
Gas chromatograms of VOCs from *T. castaneum* body extract (A) and cuticular wax (B). Bar graphs showing abundance different class of compounds detected from *T. castaneum* body extract (C) and cuticular wax (D). Peak numbers in (A) and (B) are mentioned in Table 2.

**Table 2.**
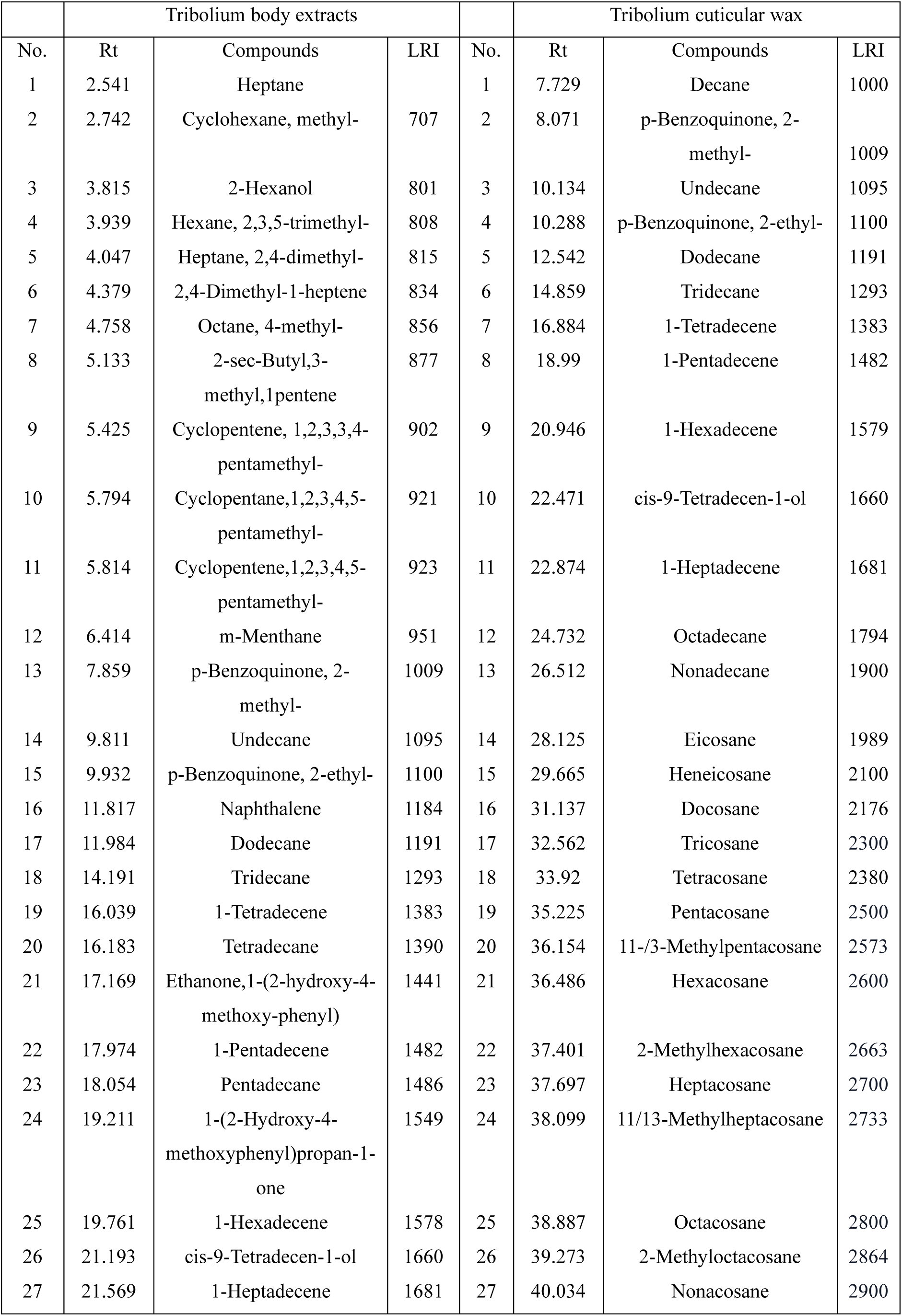
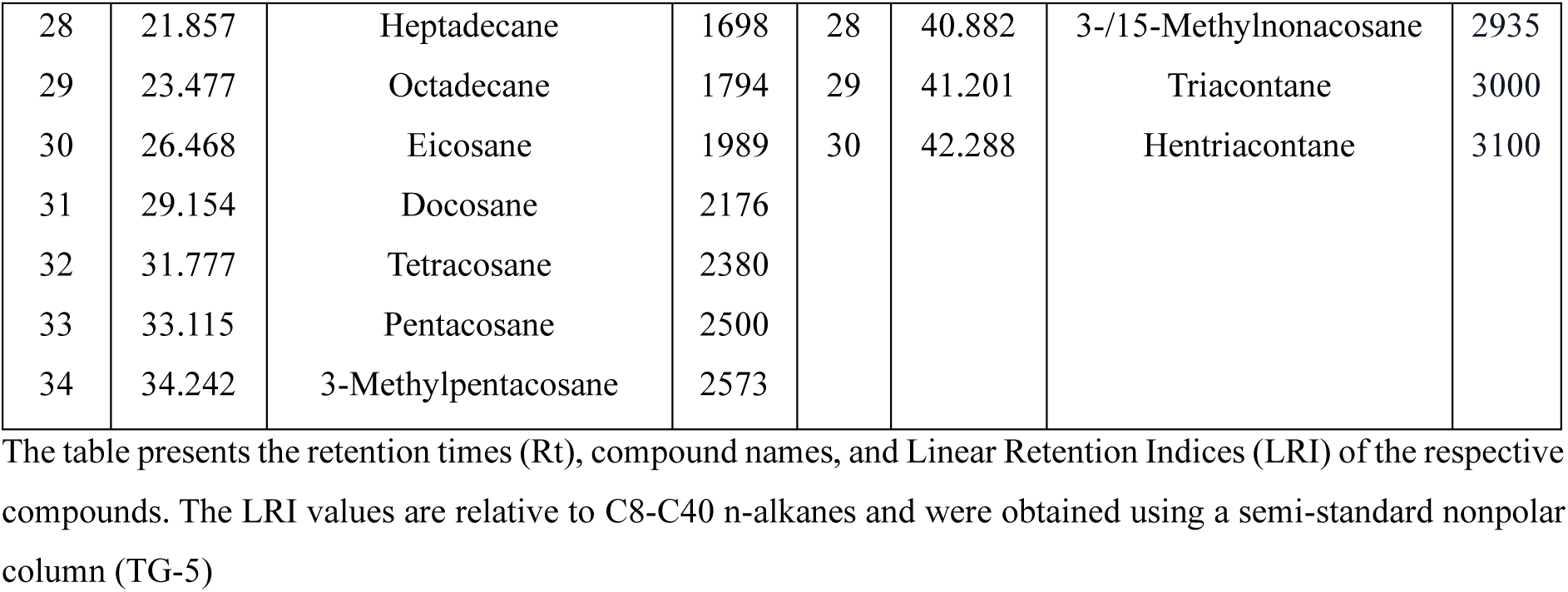
List of identified VOCs from body extracts and cuticular wax of *Tribolium castaneum*.

| Tribolium body extracts |  |  |  | Tribolium cuticular wax |  |  |  |
| --- | --- | --- | --- | --- | --- | --- | --- |
| No. | Rt | Compounds | LRI | No. | Rt | Compounds | LRI |
| 1 | 2.541 | Heptane |  | 1 | 7.729 | Decane | 1000 |
| 2 | 2.742 | Cyclohexane, methyl- | 707 | 2 | 8.071 | p-Benzoquinone, 2-methyl- | 1009 |
| 3 | 3.815 | 2-Hexanol | 801 | 3 | 10.134 | Undecane | 1095 |
| 4 | 3.939 | Hexane, 2,3,5-trimethyl- | 808 | 4 | 10.288 | p-Benzoquinone, 2-ethyl- | 1100 |
| 5 | 4.047 | Heptane, 2,4-dimethyl- | 815 | 5 | 12.542 | Dodecane | 1191 |
| 6 | 4.379 | 2,4-Dimethyl-1-heptene | 834 | 6 | 14.859 | Tridecane | 1293 |
| 7 | 4.758 | Octane, 4-methyl- | 856 | 7 | 16.884 | 1-Tetradecene | 1383 |
| 8 | 5.133 | 2-sec-Butyl,3-methyl,1pentene | 877 | 8 | 18.99 | 1-Pentadecene | 1482 |
| 9 | 5.425 | Cyclopentene, 1,2,3,3,4-pentamethyl- | 902 | 9 | 20.946 | 1-Hexadecene | 1579 |
| 10 | 5.794 | Cyclopentane,1,2,3,4,5-pentamethyl- | 921 | 10 | 22.471 | cis-9-Tetradecen-1-ol | 1660 |
| 11 | 5.814 | Cyclopentene,1,2,3,4,5-pentamethyl- | 923 | 11 | 22.874 | 1-Heptadecene | 1681 |
| 12 | 6.414 | m-Menthane | 951 | 12 | 24.732 | Octadecane | 1794 |
| 13 | 7.859 | p-Benzoquinone, 2-methyl- | 1009 | 13 | 26.512 | Nonadecane | 1900 |
| 14 | 9.811 | Undecane | 1095 | 14 | 28.125 | Eicosane | 1989 |
| 15 | 9.932 | p-Benzoquinone, 2-ethyl- | 1100 | 15 | 29.665 | Heneicosane | 2100 |
| 16 | 11.817 | Naphthalene | 1184 | 16 | 31.137 | Docosane | 2176 |
| 17 | 11.984 | Dodecane | 1191 | 17 | 32.562 | Tricosane | 2300 |
| 18 | 14.191 | Tridecane | 1293 | 18 | 33.92 | Tetracosane | 2380 |
| 19 | 16.039 | 1-Tetradecene | 1383 | 19 | 35.225 | Pentacosane | 2500 |
| 20 | 16.183 | Tetradecane | 1390 | 20 | 36.154 | 11-/3-Methylpentacosane | 2573 |
| 21 | 17.169 | Ethanone,1-(2-hydroxy-4-methoxy-phenyl) | 1441 | 21 | 36.486 | Hexacosane | 2600 |
| 22 | 17.974 | 1-Pentadecene | 1482 | 22 | 37.401 | 2-Methylhexacosane | 2663 |
| 23 | 18.054 | Pentadecane | 1486 | 23 | 37.697 | Heptacosane | 2700 |
| 24 | 19.211 | 1-(2-Hydroxy-4-methoxyphenyl)propan-1-one | 1549 | 24 | 38.099 | 11/13-Methylheptacosane | 2733 |
| 25 | 19.761 | 1-Hexadecene | 1578 | 25 | 38.887 | Octacosane | 2800 |
| 26 | 21.193 | cis-9-Tetradecen-1-ol | 1660 | 26 | 39.273 | 2-Methyloctacosane | 2864 |
| 27 | 21.569 | 1-Heptadecene | 1681 | 27 | 40.034 | Nonacosane | 2900 |
| 28 | 21.857 | Heptadecane | 1698 | 28 | 40.882 | 3-/15-Methylnonacosane | 2935 |
| 29 | 23.477 | Octadecane | 1794 | 29 | 41.201 | triacontane | 3000 |
| 30 | 26.468 | Eicosane | 1989 | 30 | 42.288 | Hentriacontane | 3100 |
| 31 | 29.154 | Docosane | 2176 |  |  |  |  |
| 32 | 31.777 | Tetracosane | 2380 |  |  |  |  |
| 33 | 33.115 | Pentacosane | 2500 |  |  |  |  |
| 34 | 34.242 | 3-Methylpentacosane | 2573 |  |  |  |  |

The table presents the retention times (Rt), compound names, and Linear Retention Indices (LRI) of the respective compounds. The LRI values are relative to C8-C40 n-alkanes and were obtained using a semi-standard nonpolar column (TG-5)

Unsaturated hydrocarbons (UHC) represented the most abundant group of odorants present in body extract (41.31 ± 5.484%) and cuticular profile (70.03 ± 8.162%), followed by Quinones (Q) [Q (body extracts) = 37.60 ± 1.677%; Q (cuticular wax) = 8.8 ± 2.308%)]. Fatty alcohol represented 12.88 ± 1.33 % and 6.79 ± 3.844 % in body extracts and the cuticular profile of T*. castaneum*, respectively. Methyl branched hydrocarbons were detected in higher amounts in cuticular wax (7.25 ± 3.939%) than body extracts (0.54 ± 0.34%) (Fig. 7C-D).

Among the 34 odorants identified from body extracts of Tribolium, 1-Pentadecene (35.16 ± 3.898 %) was most abundant followed by p-Benzoquinone, 2-ethyl-(26.08 ± 2.6 %). Apart from 1-Pentadecene, several other unsaturated hydrocarbons such as 1-Tetradecene (0.15 ± 0.051 %), 1-Hexadecene (0.34 ± 0.177 %) and 1-Heptadecene (5.66 ± 2.21 %) were detected in body extracts. cis-9-Tetradecen-1-ol (12.88 ± 1.336 %) was the only fatty alcohol detected in the body extracts of *T. castaneum*.

Cuticular wax mainly consist of various branched and unsaturated hydrocarbons as well as quinones (Table 2). 1-Pentadecene (62.07 ± 8.53 %) was the most abundant compound of cuticular wax, just like body extracts, followed by 1-Heptadecene (6.87 ± 2.615 %). Various methyl branched hydrocarbons of C25, C26, C27, C28 and C29 were detected in cuticular wax which were not detected in body extracts. Two Quinones, p-Benzoquinone, 2-methyl-(1.98 ± 0.854 %), and p-Benzoquinone, 2-ethyl-(6.81 ± 1.502 %) were also detected in cuticular wax. cis-9-Tetradecen-1-ol was also detected in considerable amount (6.79±3.844 %) in *Tribolium*’s cuticle.

Common odorants detected in both the body extracts of Tribolium and TWF included 2-Hexanol, 2-sec-Butyl-3-methyl-1-pentene, Cyclopentene, 1,2,3,3,4-pentamethyl-, Cyclopentane, 1,2,3,4,5-pentamethyl-, Cyclopentene, 1,2,3,4,5-pentamethyl-, m-Menthane, Undecane, Naphthalene, Dodecane, Tridecane, Tetradecane, 1-Pentadecene, Pentadecane, 9-Tetradecen-1-ol, 1-Heptadecene, Heptadecane and Octadecane, whereas, *Tribolium* Cuticular wax, *Tribolium* body extracts and TWF shared 7 compounds, includes Undecane, Dodecane, Tridecane, 1-Pentadecene, cis-9-Tetradecen-1-ol, 1-Heptadecene and Octadecane (Supplementary Fig. 7).

### 3.6. Effects of T. castaneum odorants in determining food preference

To determine whether the attractiveness of *T. castaneum* towards wheat flour (WF) odour increased with the addition of adult beetles’ body surface odour (cuticular wax), an additional arena bioassay was performed (Supplementary Fig. 2A). Similar to the results from our previous Y-tube assay, this experiment showed a slight increase in attraction to WF odour compared to the DCM blank, but the preference was not statistically significant (Fig. 8). While the application of *T. castaneum* body surface odour influenced the overall preference behaviour (H=20.88; df=3; p=6.88E-05), a notable number of individuals (4±1) remained indecisive, a phenomenon not observed in earlier arena bioassays (Fig. 8). Interestingly, *T. castaneum* tended to avoid the WF odour enriched with beetles’ body surface odour (1±1), showing a preference for the fresh flour odour instead (3±1).

**Fig. 8.**
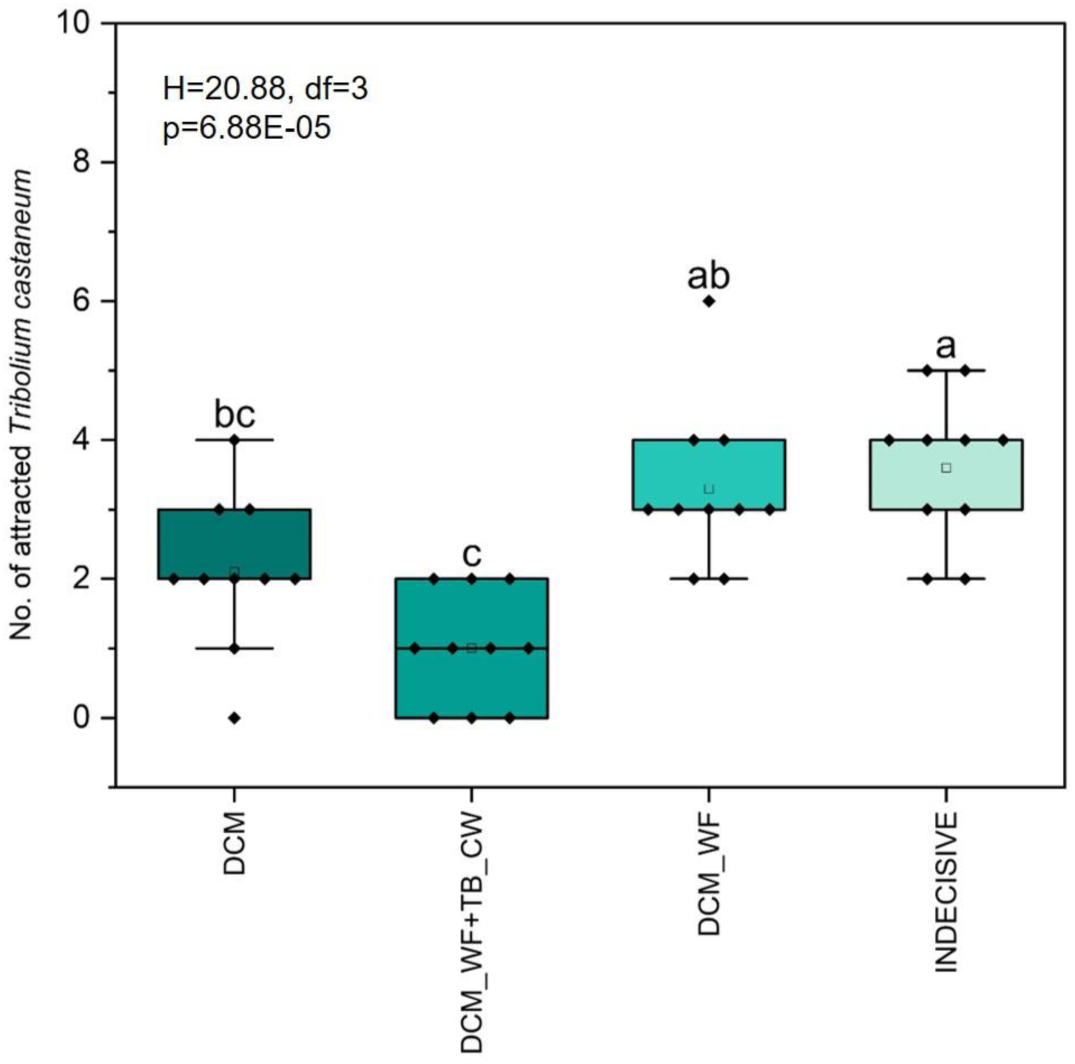
Box plot representing the attractiveness of adult *T. castaneum* towards DCM extract of wheat flour (WF: 20 µl) and DCM extract of wheat flour (20 µl) mixed with cuticular wax extract of *T. castaneum* (10 µl). Statistical significance (p < 0.05) was determined using the Kruskal-Wallis test followed by a Dunn’s post-hoc multiple comparison test. Letters above the box plots indicate statistical differences, where box plots sharing the same letter show no significant difference in beetle attraction.

### 3.7. Role of infested flour odour with different concentrations of 1-Pentadecene on T. castaneum preference behaviour

In our study (Supplementary Fig. 2B), we observed that wheat flour (WF) enriched with *T. castaneum* body odour did not attract the beetles. However, in a previous Y-tube bioassay (Fig. 4), odours from *T. castaneum*-infested flour significantly attracted more beetles.

To further investigate whether the concentration of specific odour compounds in *T. castaneum*-infested flour influences beetles’ behavioural response, we analyzed wheat flour samples with varying *T. castaneum* infestation levels. Odour profile analysis showed that the contents of 1-Pentadecene varied considerably among these samples. We chose infested flour samples whose 1-Pentadecene contents were 10 ng/g, 20 ng/g and 30 ng/g, extracted their odours in DCM and tested their attractiveness to beetles against a DCM blank. The results showed that *T. castaneum* preference for flour was decreased as the concentration of 1-Pentadecene was increased although this trend was not statistically significant (H=5.728; df=3; p=0.1256) (Fig. 9).

**Fig. 9.**
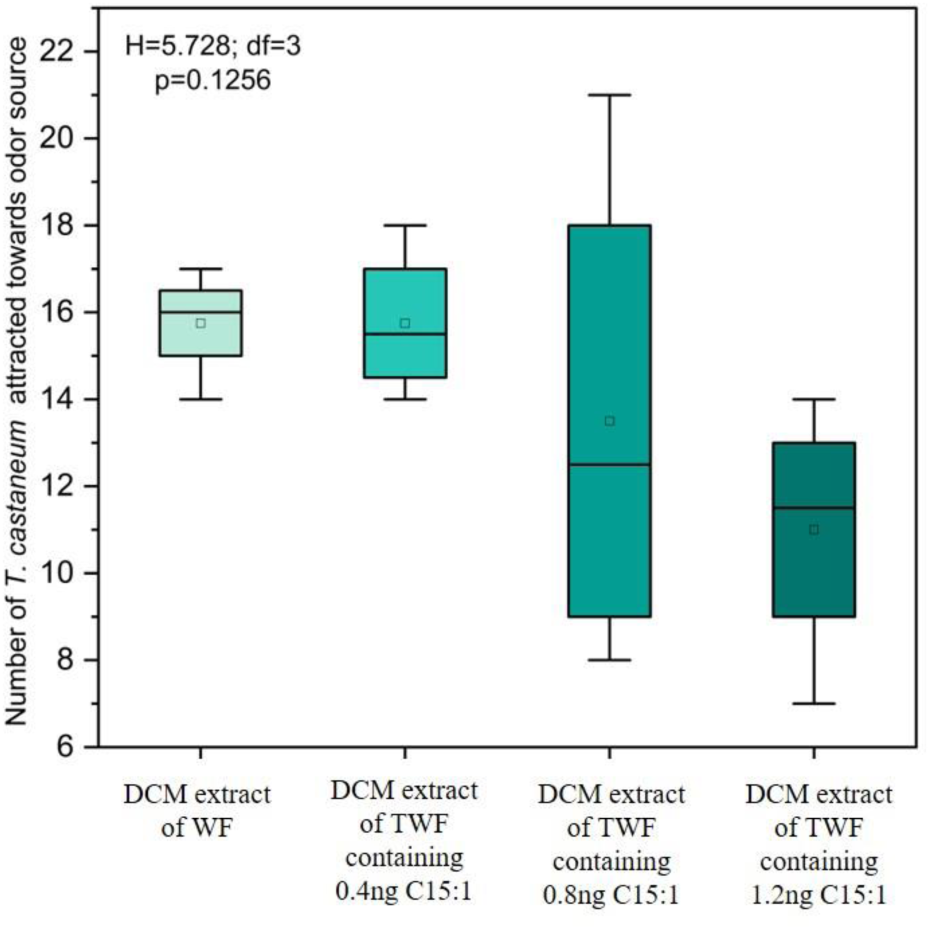
Box plot representing the number of adult *T. castaneum* attracted to infested wheat flour odour (20µl DCM extract) containing 0.4 ng, 0.8 ng, and 1.2 ng of 1-Pentadecene compared to control fresh flour (0 ng 1-Pentadecene). Statistical significance (p < 0.05) was determined using the Kruskal-Wallis test.

### 3.8. Effects of T. castaneum odorants in determining reproductive success

An experiment was designed to determine the reproductive success of *T. castaneum* using different concentrations of 1-Pentadecene in extracts. Three experimental sets were prepared, each containing 15 g of TWF containing 10 ng/g, 20 ng/g, and 30 ng/g of pentadecane respectively, and inoculated with 10 adult beetles (♂5: ♀5). After 90 days, the number of adult *T. castaneum* in each set was counted (Supplementary Fig. 3). The results showed a decreasing trend in reproductive success as the concentration of 1-Pentadecene increased (Fig. 10). The highest number of adults (∼140 ) were produced with the lowest concentration of 10 ng/g, whereas higher concentrations of 20 ng/g and 30 ng/g resulted in even less, ∼90 and ∼60 adults, respectively. Although this decline trend may indicate that higher levels of 1-Pentadecene in extracts could suppress reproductive success, this effect is not statistically significant (p = 0.0594) (Fig.10).

**Fig. 10.**
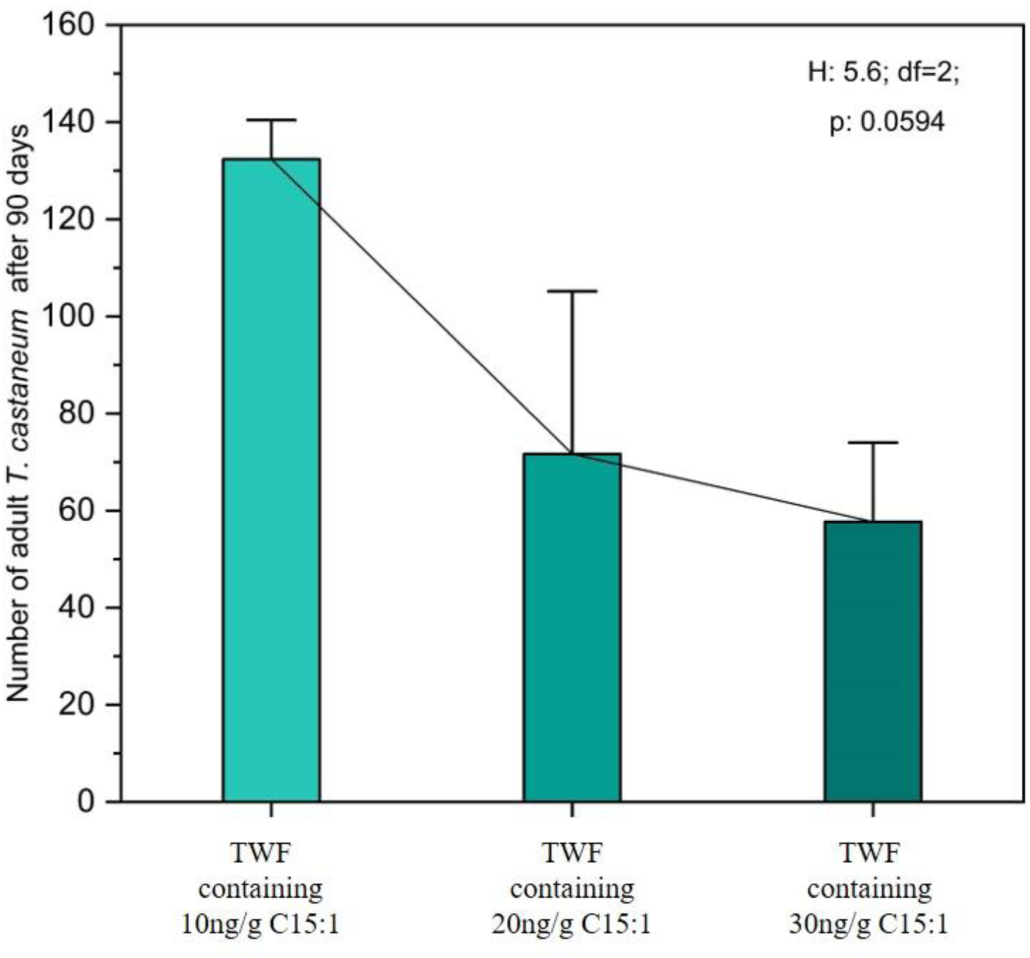
Bar plot representing the number of adult *T. castaneum* after 90 days post-inoculation (♂5: ♀5) in three experimental sets. Each set contained 15 g of wheat flour supplemented with *T. castaneum*-infested wheat flour (TWF) DCM extract. The concentration of 1-Pentadecene was maintained at 10 ng/g, 20 ng/g, and 30 ng/g in the respective experimental sets. Statistical significance (p<0.05) was measured using Kruskal Wallis test.

### 3.9. Molecular docking analysis of important odorants with T. castaneum olfactory receptor (TcOR1)

In silico molecular docking study, was conducted to evaluate the binding potential of two key odorants with the *T. castaneum* odorant receptor, TcOR1. Initially, the aggregation pheromone 4,8-DMD was selected for blind docking with TcOR1 to identify potential binding pocket formed between the receptor and the odorant. Once the binding site was predicted, 1-Pentadecene was subsequently docked into the same pocket where 4,8-DMD had previously shown affinity, to assess its binding interactions with TcOR1. The grid box dimension values are in Supplementary Table 5. The binding affinity *(ΔG)* of TcOR1 for both odorants was found to be -5.8 kcal/mol for 4,8-DMD and -5.7 kcal/mol for 1-Pentadecene, indicating a similar high strength of the interaction between TcOR1 and both odorants (Table 3; Fig. 11). Key residues that participated in the binding sites of TcOR1 to both odorants were MET49, PHE69, THR72, GLN73, THR134, VAL135, TYR138, LEU154, ILE177, GLY180, ALA181, and TYR286. Interestingly, in the case of 1-Pentadecene, another binding interaction involving ASN184 was observed. Predictions calculated for both odorants involved alkyl, pi-alkyl, and van der Waals forces. The scores also show that with 4,8-DMD, there was a predicted unfavourable bump in the docking but that this also could represent possible steric hindrance or poorer interacting capability within the pocket (Table 3; Fig. 11). Another two odorants of *Tribolium,* p-Benzoquinone-2-methyl-, and p-Benzoquinone-2-ethyl-, were also docked with the same pocket of TcOR1 to check their binding potential with that odorant receptor. Docking results showed considerable strong binding affinity (TcOR1-p-Benzoquinone-2-methyl - complex ΔG=-5.0 and TcOR1-p-Benzoquinone-2-ethyl-complex ΔG= -5.0) (Table 3; Supplementary Fig. 8).

**Fig. 11.**
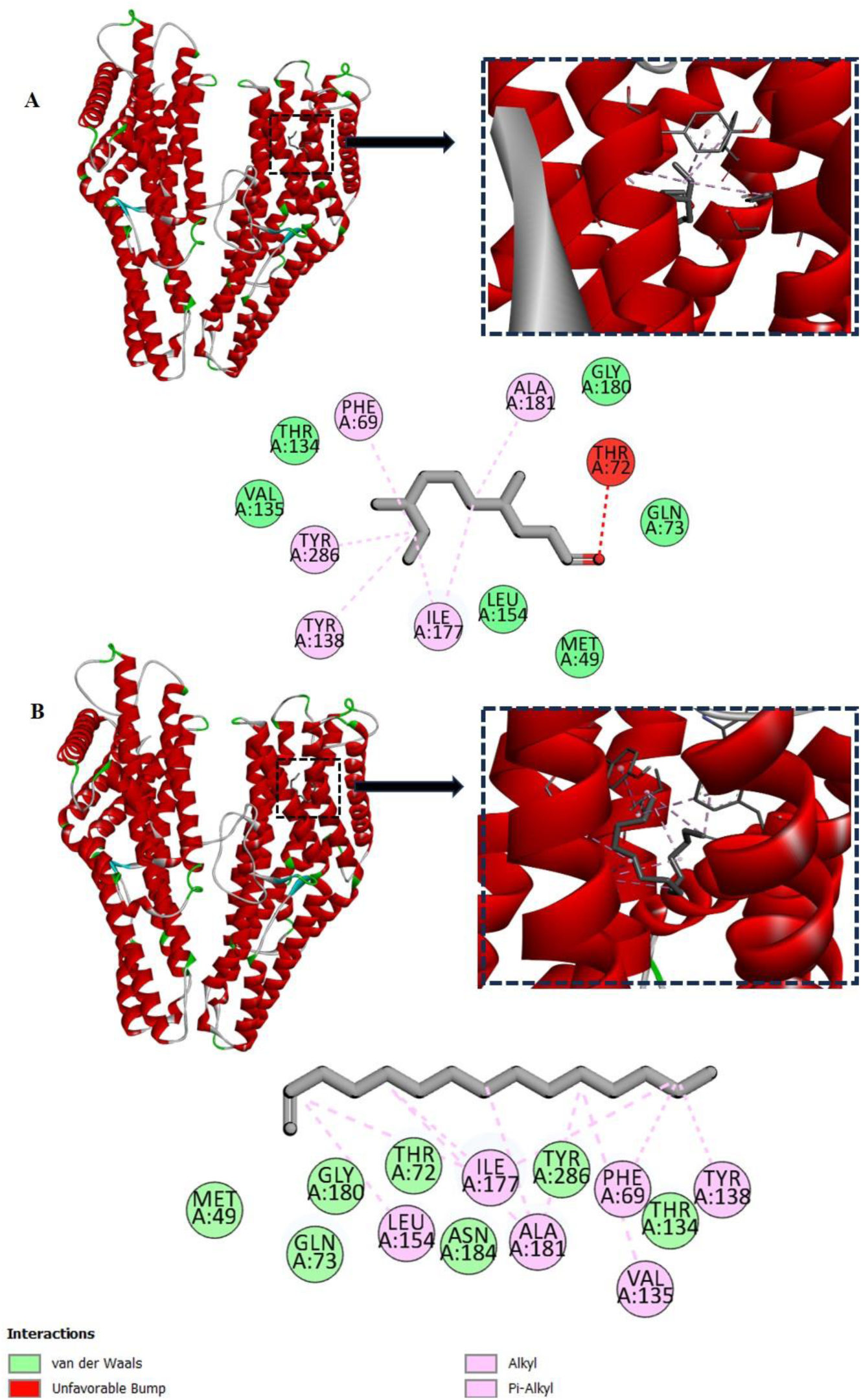
*In-silico* molecular docking visualization of TcOR1 (odorant receptor) complexed with 4,8-DMD (A) and 1-Pentadecene (B), using Discovery Studio Visualizer v21.1.0.20298. The 3D protein-ligand interaction maps are shown with the receptor in red helical ribbons. Ligand-binding details are displayed in close-up insets, along with 2D diagrams highlighting key residues involved in interactions.

**Table 3.**
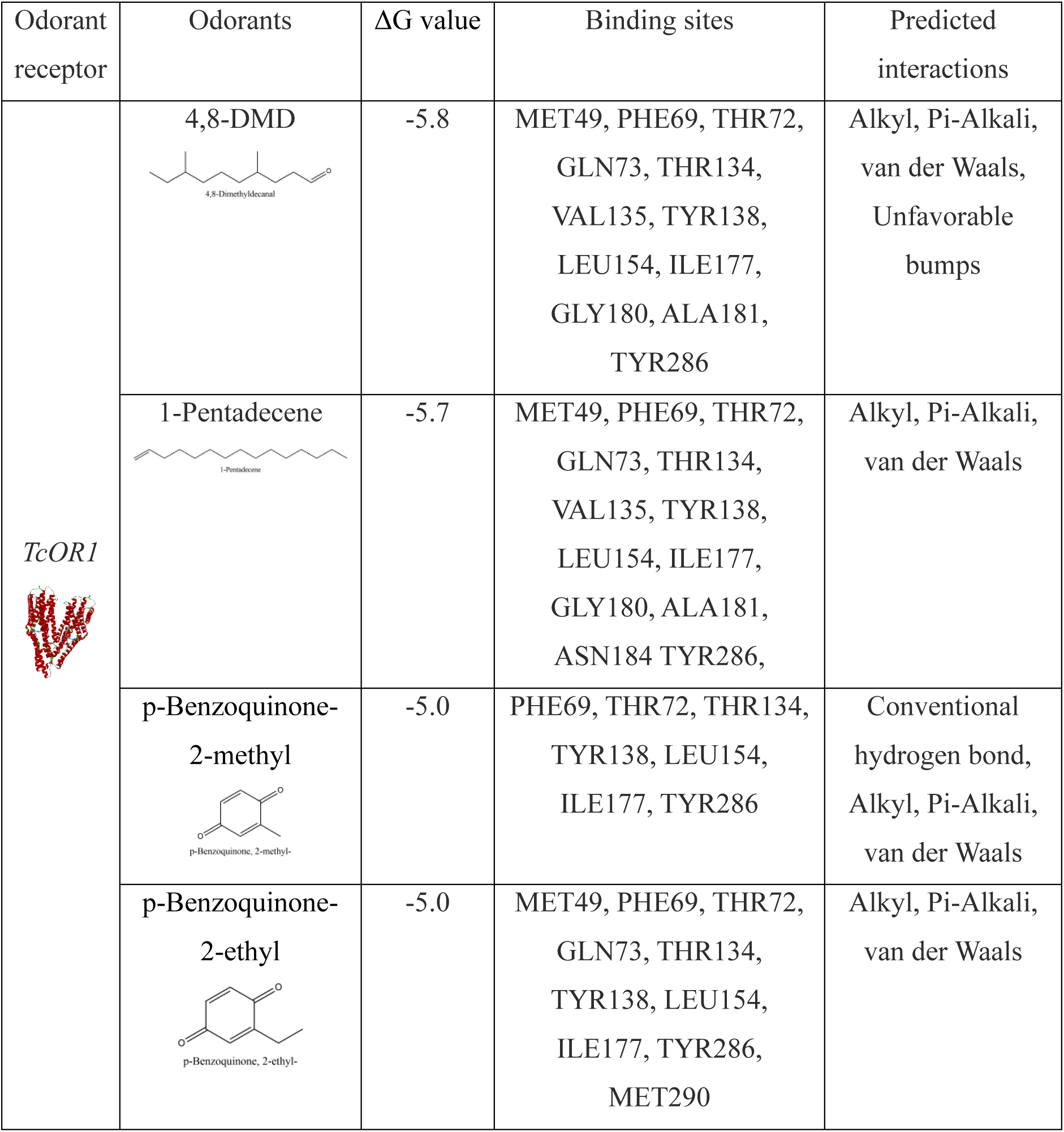
*In-silico* result showing binding potential (ΔG) and type of interactions of various odorants with TcOR1 of *T. castaneum*.

## 4. Discussion

### 4.1. Alternation of nutritional profiles in response to flour beetle infestation

The present investigation revealed that *T. castaneum* infestation significantly altered the nutrient profile of wheat flour, including a reduction in sugar and lipid content and an increase in protein content. Previous studies have reported that *T. castaneum* prefers to feed on wheat germ and endosperm over wheat bran and can selectively consume the germ and endosperm when all three components are mixed (Waldbauer and Bhattacharya, 1973). This preference results in the consumption of a larger portion of the endosperm, which constitutes 80–85% of wheat flour and is the primary source of starch and sugars (Dale and Housley, 1986; Fardet, 2010).Wheat germ, although only 3% of whole wheat, is also rich in carbohydrates. However, the flour beetle does not highly prefer wheat bran, which makes up approximately 18% of wheat and has a relatively low sugar content (Fardet, 2010; Pomeranz, 1988). Therefore, the observed reduction in sugar content in *T. castaneum*-infested wheat flour can be attributed to the insect’s feeding preferences to wheat endosperm and germ over bran.

Although the total lipid content decreased, this reduction was not statistically significant. While wheat germ contains a high amount of lipids, the endosperm has relatively low lipid levels, and both are preferred by *T. castaneum*. In contrast, wheat bran, which is less preferred by the insect, contains a considerable amount of lipids, possibly explaining the non-significant reduction in lipid content in the infested wheat flour. Further evidence supporting the reduction of lipids due to red flour beetle’s exploitation of germ and endosperm comes from the differential quantities of oleic and linoleic acids between fresh wheat flour (WF) and infested wheat flour (TWF). Durate et al. (2021) reported that these two fatty acids comprise more than 50% of the total fatty acids in *T. castaneum* larvae, pupae, and adults, as they are essential for the insect’s development (Duarte et al., 2021). Wheat germ is particularly rich in these fatty acids compared to bran, while the endosperm contains negligible amounts (Geng et al., 2015). In this study, the significant decrease in unsaturated fatty acids, particularly linoleic acid, in TWF suggests that the change in fatty acid content is likely due to the insect’s selective feeding on specific portions of the wheat. During the experiment, the flour beetle likely consumed a substantial amount of wheat germ, which may have contributed to the reduced relative abundance of unsaturated fatty acids in the infested flour.

In addition, breakdown of storage lipids (TAG) and an increase in FFA content were observed during *T. castaneum* infestation. Free fatty acids are more prone to undergo hydrogenation and oxidation that the esterified fatty acids (Tena et al., 2019), leading to rancidity and the formation of new lipophilic odorant molecules (Aher et al., 2022). As a result, the nutritional quality of the flour is compromised, or the development of off-flavours reduces its acceptability and commercial value.

Interestingly, total soluble protein content slightly increased in TWF, possibly due to the presence of insect tissue, eggs, and debris in the flour, a finding consistent with the observations of Carvalho et al. (2023) in *T. castaneum*-infested wheat flour (Carvalho et al., 2023). Additionally, the content of secondary metabolites, such as phenolics, flavonoids, and terpenoids, increased in TWF. This increase is likely due to the insect’s exploitation of the endosperm and germ, which contain lower levels of secondary metabolites, while residual bran parts—rich in phenolic acids, polyphenols, and other compounds-remain (Fardet, 2010).

### 4.2. *Behavioural alternation of T. castaneum* in response to WF and TWF

The odour of wheat flour has been reported to influence the food preference of *T. castaneum* (Gao et al., 2024; Seifelnasr et al., 1982), with beetles showing a distinct preference for wheat flour that has already been infested by *T. castaneum* (Stevenson et al., 2017). In the present investigation, using both a two-choice Y-tube bioassay and an arena bioassay, it was found that the DCM extract of uninfested wheat flour did not attract adult *T. castaneum*. However, the DCM extract of *T. castaneum*-infested wheat flour attracted a significant number of *T. castaneum* adults, both when compared to a blank control and to the DCM extract of uninfested wheat flour. This result aligns with the findings of Stevenson et al. (2017), who observed a similar response using fresh wheat flour and *T. castaneum-*infested flour rather than DCM extracts. The present study also revealed that methanolic extracts from the same fractions did not significantly affect the food preference of *T. castaneum*, suggesting that nonpolar compounds soluble in DCM play a key role in influencing the beetle’s food preference, whereas polar compounds soluble in methanol do not.

In a separate behavioural assay, the adult *T. castaneum* was provided with 24 hours to infest either fresh wheat flour or pre-infested nutrition depleted flour. Beetles preferred fresh wheat flour, but no statistically significant difference was found. This preference may even go further than chemical cues such as the quality of diets or even texture of the flour, which can be ascribed by food selection behaviour of the species (Astuti et al., 2020). The colour of *T. castaneum*-infested wheat flour (dark) compared to fresh wheat flour (light) may also influence the behaviour of adult beetles. Wagner et al. (2018) found that *T. castaneum* adults exhibit a preference for lighter-coloured food over darker-coloured food (Wagner et al., 2018). Thus, it can be assumed that while *T. castaneum* adults may initially be drawn to infested flour based on chemical cues, as demonstrated by the Y-tube olfactometer and arena bioassays, but over a longer duration, they tend to settle in flour that is nutritionally superior or possesses other desirable physical characteristics. However, in a separate experimental setup, adults of *T. castaneum* preferred to colonise the wheat flour that was enriched with the DCM extract from *T. castaneum-infested flour* over fresh wheat flour over a 24-hour span. It indicates that chemical signals from *T. castaneum*-infested flour might improve the appeal of fresh flour to adult red flour beetles, increasing their likelihood of preferring and colonising flour that has been exposed to these chemical cues.

### 4.3. Altered odour profile of T. castaneum-infested wheat flour

Analysis of the DCM extracts from both fresh and *T. castaneum* infested wheat flour revealed a total of 46 volatile compounds. Among these, 26 volatiles were common in both tested flours, constituting 87.79 ± 0.938 % of the total volatiles detected in fresh wheat flour and 70.96 ± 0.718% in *T. castaneum*-infested wheat flour. No significant differences were observed in the relative abundances of these common compounds. A similar finding was reported by Niu et al. (2016), who demonstrated that 21 volatile compounds shared between fresh and *T. castaneum*-infested wheat flour exhibited no change in quantity, suggesting that *T. castaneum* infestation does not considerably alter the pre-existing volatile profile of wheat flour. However, our study identified certain volatile compounds that were contributed by *T. castaneum,* during infestation. Specifically, 10 such volatiles were detected exclusively in the DCM extract of *T. castaneum*-infested wheat flour, including 1-Pentadecene, cis-9-Tetradecen-1-ol, 1-Heptadecene, and 4,8-Dimethyldecanal (4,8-DMD). Principal component analysis (PCA) identified these compounds as key chemomarkers for *T. castaneum*-infested wheat flour.

Notably, 4,8-DMD, a fatty acid derivative, is produced by the male *T. castaneum* and has been reported to attract both sexes of the species (Kim et al., 2005; Ming and Lewis, 2010; Rath et al., 2021). Additionally, 1-Pentadecene, another important species-specific hydrocarbon, was also detected in significant quantities in *T. castaneum*-infested flour. This volatile has been recognized as a chemomarker for *T. castaneum* infestation by multiple authors (Đukić et al., 2021; Tian et al., 2022; Villaverde et al., 2007) and plays a dual role in regulating the beetle’s behaviour (Đukić et al., 2021). Đukić et al. (2021) reported that at low concentrations, 1-Pentadecene acts as an attractant for adult *T. castaneum*, while at higher concentrations, it functions as a repellent.

The presence of these two compounds was further validated using HS-SPME-GCMS, further validating their importance as chemomarkers for *T. castaneum* infestation (Supplementary Table 2 ). Additionally, several other compounds detected exclusively in *T. castaneum*-infested flour, except 4,8-DMD (genus specific aggregation pheromone), have been reported to exhibit semiochemical properties in other species of Coleopteran beetles (Supplementary Table 3).

While investigating the sources of these volatiles detected in *Tribolium*-infested wheat flour, it was found that some compounds, such as 1-Pentadecene, 1-Heptadecene, and cis-9-Tetradecen-1-ol, are produced by *T. castaneum*, which eventually secreted into the flour. These compounds have been detected in the cuticular wax and whole-body extract of *T. castaneum* during our study, suggesting that the transmission of these compounds to the wheat flour may occur through direct physical contact between the beetles and the flour. Interestingly, 4,8-Dimethyldecanal (4,8-DMD), although not detected in either body extracts or the cuticular wax of *T. castaneum*, has been well-documented as a volatile compound produced by the species. This discrepancy might be explained by the possibility that 4,8-DMD is not constitutively produced but rather synthesized by the beetles at specific times to perform certain behavioural functions.

On the other hand, two major defensive chemicals of *T. castaneum*, p-Benzoquinone, 2-methyl- and p-Benzoquinone, 2-ethyl-, were not detected in the infested wheat flour, neither in the DCM extract nor via SPME-guided sampling. However, previous reports suggest the occurrence of p-Benzoquinone, 2-methyl-in *T. castaneum-infested flour* (Niu et al., 2016). Another compound, Acetophenone, was detected in *T. castaneum* infested wheat flour, although it was not found in either the cuticular wax or whole-body extracts of *T. castaneum*. However, Villaverde et al. (2007) identified Acetophenone in very small quantities from *T. castaneum* using SPME-guided GC-MS with a CAR/PDMS fiber. This suggests that *T. castaneum* may release Acetophenone in very low amounts over time, which eventually accumulate in the flour and eventually be detected. Additionally, Acetophenone is known to be a breakdown product of benzenoid volatiles, such as Ethylbenzene and Xylene (Azeez et al., 2023), which were slightly reduced in infested flour compared to fresh flour, during our investigation.

Several other volatiles detected in the infested flour were not directly linked to *T. castaneum*, including Neral, 2-Heptanone, 2,3,3-Trimethyl-1-hexene, and Cyclohexanol, 1-[3-(1-pyrrolidinyl)-1-propynyl]. Among these, 2-Heptanone is reported to be produced from oxidation of lipids, particularly from fatty acids, during long-term storage (Xi et al., 2023). According to Balakrishnan et al. (2017), 2-Heptanone elicits a strong antennal response in both sexes of *T. castaneum*, even at low concentrations, and the beetles’ antennae show an even stronger response to this molecule than to some major compounds like 4,8-DMD, 1-Pentadecene, and Quinones. In addition to 2-heptanone, Citral (neral) was also tested by the authors, revealing that Citral can evoke a strong antennal response, particularly in male *T. castaneum* (Balakrishnan et al., 2017).

All of the aforementioned compounds that could modulate the behaviour of *T. castaneum* were detected only in the DCM extract of *T. castaneum* infested wheat flour, but were absent in the methanolic extract, as shown in Fig.6. This likely explains why, during the behavioural assays where the odours of fresh and *T. castaneum* infested wheat flour were compared, the DCM extracts from the infested flour elicited significant responses, while the methanolic extracts from the same infested flour did not.

### 4.4. Differed behavioural response of T. castaneum towards TWF DCM extract and WF DCM extract combined with cuticular wax of T. castaneum

The present investigation revealed that although the volatile blend from *T. castaneum*-infested wheat flour attracted adult beetles, the combination of volatiles from fresh flour and cuticular wax of *T. castaneum* failed to do so in an arena bioassay. Instead, it repelled the tested individuals. This suggests that certain compounds present in the cuticular wax but absent from the infested wheat flour acted as repellents, making the odour mixture undesirable to the adults. Two such compounds, p-benzoquinone, 2-methyl-, and p-benzoquinone, 2-ethyl-, are found in significant quantities in the cuticular wax fraction (1.98 ± 0.854 % and 6.81 ± 1.502 %, respectively) but are entirely absent from infested wheat flour. These compounds are known to be defence-related chemicals, produced when the flour becomes overcrowded, and serve as repellents to prevent further infestation. This is an adaptive strategy to avoid overcrowded environments. Additionally, literature suggests that high concentrations of these volatiles may even be lethal to the species. This likely explains why adult beetles preferred the volatiles from infested wheat flour, which contained 4,8-DMD and lacked quinones, over the combination of fresh flour volatiles and *T. castaneum* cuticular wax, which contained significant amounts of Benzoquinones.

In addition to Benzoquinones, 1-Pentadecene is another component that showed varying relative abundances between *T. castaneum*-infested wheat flour extract (amount of 1-Pentadecene = 12.09 ± 0.198%) and fresh flour extract supplemented with beetles ’cuticular wax (amount of 1-Pentadecene in cuticular wax = 62.07 ± 8.53%). The differential behavioural response of *T. castaneum* towards infested flour extract and the combination of fresh flour extract and *T. castaneum* cuticular wax can be explained by the work of Đukić et al. (2021). In their study, Đukić et al. (2021) demonstrated that wheat bran with low concentrations of 1-Pentadecene is more attractive to *T. castaneum*, while slightly higher concentrations have a neutral effect. However, at the highest concentration, 1-Pentadecene exhibited a strong repellent effect on adult *T. castaneum*. To further support our findings, we tested the attractiveness of different concentrations of 1-Pentadecene on *T. castaneum* using a Y-tube olfactometer. The results mirrored those of Đukić et al. (2021), showing that with increasing doses of 1-Pentadecene, *T. castaneum’*s preference for the odour source gradually declined, although the change was not statistically significant.

### 4.5. Putative growth regulatory activity of specific T. castaneum odour

The present study also revealed that an increase in the concentration of 1-Pentadecene in wheat flour appears to reduce the reproductive success of *T. castaneum*. This may be due to the interaction of multiple volatile compounds. Previous reports suggest that 4,8-DMD, the primary aggregation pheromone, promotes *T. castaneum* aggregation and subsequently influences mating success. On the other hand, 1-Pentadecene can act either as an attractant or a repellent depending on its concentration (Đukić et al., 2021). As both 4,8-DMD and 1-Pentadecene are lipid derivatives, their biosynthesis may be regulated by substrate availability (Kim et al., 2005; Ney and Boland, 1987). According to (Duehl et al., 2011), the release of Benzoquinones is correlated with 1-Pentadecene concentration, and Benzoquinones have been shown to inhibit the biosynthesis of 4,8-DMD by interfering with lipid accumulation. Additionally, Benzoquinones promote lipid desaturation, which facilitates 1-Pentadecene production (Wang, 1981). It is therefore possible that 1-Pentadecene serves as a regulator of population growth for *T. castaneum* in confined environments. Initially, 1-Pentadecene and 4,8-DMD work together to increase the population, but as the population grows, 1-Pentadecene, through its influence on benzoquinone production, decreases 4,8-DMD levels, leading to a decline in aggregation and eventually stabilising the population.

Furthermore, the present study also suggests that both 1-Pentadecene and Benzoquinones can interact with the odorant binding site of TcOR1, which is also the receptor for 4,8-DMD. When 1-Pentadecene or Benzoquinones bind to TcOR1, they may reduce the receptor’s sensitivity to 4,8-DMD. Previous studies have shown that malfunction of TcOR1, where 4,8-DMD cannot interact with the receptor, affects behavioural responses of *T. castaneum* and reduces reproductive success. It is possible that at low concentrations of 1-Pentadecene, 4,8-DMD binds to TcOR1 and triggers aggregation behaviour. However, at higher concentrations of 1-Pentadecene, the compound may competitively bind to TcOR1, preventing *T. castaneum* from detecting the aggregation cues, thereby inhibiting aggregation. It can be assumed that 1-Pentadecene plays a key role in regulating the population size of the red flour beetle by influencing aggregation behaviour. Either by interfering with the detection of the primary aggregation pheromone, 4,8-DMD, or by inhibiting the production of 4,8-DMD and ultimately impacting the reproductive success and population size of the species.

## 5. Conclusion

this study highlights the significant impact of *T. castaneum* infestation on the nutritional composition of stored wheat flour and reveals key chemical interactions influencing beetle behaviour. The reduction in sugar and lipid content, particularly linoleic acid, indicates a substantial decline in nutritional quality. Additionally, the increase in phenolic and tannin contents, along with the accumulation of beetle-derived odours, reduces the digestibility of the flour and creates a malodour, diminishing the value of infested flour and that of the products made from it. This study also identifies certain chemical markers, such as 1-Pentadecene, cis-9-Tetradecen-1-ol, and 1-Heptadecene, associated with *T. castaneum* infestation. The early detection of these markers could enhance the security of stored grains. Furthermore, multiple volatiles were found to modulate *T. castaneum* behaviour, notably 4,8-Dimethyldecanal (4,8-DMD) and 1-Pentadecene. Wheat flour containing similar amounts of 4,8-DMD but higher concentrations of 1-Pentadecene was avoided by adult beetles, suggesting that elevated levels of 1-Pentadecene may suppress the aggregation response typically mediated by 4,8-DMD. This was validated through molecular docking studies, which explored the interaction of these two odorants with the TcOR1 odorant receptor of *T. castaneum*. The lack of aggregation response led to reduced population growth in the presence of higher concentrations of 1-pentadecene. However, further research is needed to address few questions, including whether the population size of *T. castaneum* influences the production of 1-Pentadecene in individual beetles, which is subsequently transmitted to the flour and may help regulate population size. Additionally, understanding the specific ratio of 4,8-DMD and 1-Pentadecene at which 1-Pentadecene suppresses the effects of 4,8-DMD remains a key area for future investigation.

## Acknowledgements

This research was conducted at the Semiochemicals and Lipid Laboratory, Department of Life Sciences, Presidency University, Kolkata. We extend our heartfelt gratitude to the head of this lab, Prof. Mousumi Poddar Sarkar, for her invaluable guidance and support. We also acknowledge the DBT-BUILDER programme, Government of India (BT/INF/22/SP45088/2022, dated 17.02.2022), for providing the GC-MS facilities at the Department of Life Sciences, Presidency University, Kolkata. Special thanks are extended to Dr. Ritwika Bera and Pritom Das for their generous cooperation and assistance throughout the course of this work.

## CRediT authorship contribution statement

**Subhadeep Das**: Conceptualization, Investigation, Methodology, Formal analysis, Writing-Original draft preparation, Data curation, Writing-Reviewing and Editing. **Sourav Manna**: Conceptualization, Visualization, Writing, Methodology, Validation. **Oishika Chatterjee**: Data curation, Reviewing, Methodology. **Riya Saha**: Data curation, Reviewing, Methodology. **Oishee Janet Sarkar**: Data curation, Reviewing, Methodology

## Conflict of interest

All authors have seen and approved the manuscript, and it hasn’t been accepted or published elsewhere. The authors declare that they have no conflict of interest.

## Funding sources

This research did not receive any specific grant from funding agencies in the public, commercial, or not-for-profit sectors.

## Notes

### Competing Interest Statement

The authors have declared no competing interest.

## References

Aher, R.R., Reddy, P.S., Bhunia, R.K., Flyckt, K.S., Shankhapal, A.R., Ojha, R., Everard, J.D., Wayne, L.L., Ruddy, B.M., Deonovic, B., Gupta, S.K., Sharma, K.K., Bhatnagar-Mathur, P., 2022. Loss-of-function of triacylglycerol lipases are associated with low flour rancidity in pearl millet [Pennisetum glaucum (L.) R. Br.]. Front. Plant Sci. 13. 10.3389/fpls.2022.962667

Ai, S., Zhang, Y., Chen, Y., Zhang, T., Zhong, G., Yi, X., 2022. Insect-Microorganism Interaction Has Implicates on Insect Olfactory Systems. Insects 13, 1094. 10.3390/insects13121094

Ainsworth, E.A., Gillespie, K.M., 2007. Estimation of total phenolic content and other oxidation substrates in plant tissues using Folin–Ciocalteu reagent. Nat. Protoc. 2, 875–877. 10.1038/nprot.2007.102

Astuti, L.P., Rizali, A., Firnanda, R., Widjayanti, T., 2020. Physical and chemical properties of flour products affect the development of *Tribolium castaneum*. J. Stored Prod. Res. 86, 101555. 10.1016/j.jspr.2019.101555

Athanassiou, C.G., Kavallieratos, N.G., Campbell, J.F., 2016. Capture of Tribolium castaneum and Tribolium confusum (Coleoptera: Tenebrionidae) in Floor Traps: The Effect of Previous Captures. J. Econ. Entomol. 109, 461–466. 10.1093/jee/tov307

Atta, B., Rizwan, M., Sabir, A.M., Gogi, M.D., Ali, K., 2020. Damage potential of Tribolium castaneum (Herbst) (Coleoptera: Tenebrionidae) on wheat grains stored in hermetic and non-hermetic storage bags. Int. J. Trop. Insect Sci. 40, 27–37. 10.1007/s42690-019-00047-0

Azeez, M.O., Nafiu, S.A., Olarewaju, T.A., Olabintan, A.B., Tanimu, A., Gambo, Y., Aitani, A., 2023. Selective Catalytic Oxidation of Ethylbenzene to Acetophenone: A Review of Catalyst Systems and Reaction Mechanisms. Ind. Eng. Chem. Res. 62, 12795– 12828. 10.1021/acs.iecr.3c01588

Balakrishnan, K., Holighaus, G., Weißbecker, B., Schütz, S., 2017. Electroantennographic responses of red flour beetle Tribolium castaneum Herbst (Coleoptera: Tenebrionidae) to volatile organic compounds. J. Appl. Entomol. 141, 477–486. 10.1111/jen.12366

Bligh, E.G., Dyer, W.J., 1959. A rapid method of total lipid extraction and purification. Can. J. Biochem. Physiol. 37, 911–917. 10.1139/o59-099

Bradford, M.M., 1976. A rapid and sensitive method for the quantitation of microgram quantities of protein utilizing the principle of protein-dye binding. Anal. Biochem. 72, 248–254. 10.1016/0003-2697(76)90527-3

Campbell, J.F., 2012. Attraction of walking *Tribolium castaneum* adults to traps. J. Stored Prod. Res. 51, 11–22. 10.1016/j.jspr.2012.06.002

Carvalho, M.O., Geirinhas, H., Duarte, S., Graça, C., de Sousa, I., 2023. Impact of red flour beetle infestations in wheat flour and their effects on dough and bread physical, chemical, and color properties. J. Stored Prod. Res. 102, 102095. 10.1016/j.jspr.2023.102095

Christie, W., 1993. PREPARATION OF ESTER DERIVATIVES OF FATTY ACIDS FOR CHROMATOGRAPHIC ANALYSIS.

Csepregi, K., Kocsis, M., Hideg, É., 2013. On the spectrophotometric determination of total phenolic and flavonoid contents. 10.1556/abiol.64.2013.4.10

Dale, E.M., Housley, T.L., 1986. Sucrose synthase activity in developing wheat endosperms differing in maximum weight. Plant Physiol. 82, 7–10. 10.1104/pp.82.1.7

Dissanayaka, D.M.S.K., Sammani, A.M.P., Wijayaratne, L.K.W., 2020. Orientation of *Tribolium castaneum* (Herbst) (Coleoptera: Tenebrionidae) adults at various distances to different concentrations of aggregation pheromone 4,8-dimethyldecanal. J. Stored Prod. Res. 87, 101631. 10.1016/j.jspr.2020.101631

Dissanayaka, D.M.S.K., Sammani, A.M.P., Wijayaratne, L.K.W., 2018. Aggregation pheromone 4,8-dimethyldecanal and kairomone affect the orientation of *Tribolium castaneum* (Herbst) (Coleoptera: Tenebrionidae) adults. J. Stored Prod. Res. 79, 144– 149. 10.1016/j.jspr.2018.07.005

Duarte, S., Limão, J., Barros, G., Bandarra, N.M., Roseiro, L.C., Gonçalves, H., Martins, L.L., Mourato, M.P., Carvalho, M.O., 2021. Nutritional and chemical composition of different life stages of *Tribolium castaneum* (Herbst). J. Stored Prod. Res. 93, 101826. 10.1016/j.jspr.2021.101826

Duehl, A.J., Arbogast, R.T., Teal, P.E.A., 2011. Density-Related Volatile Emissions and Responses in the Red Flour Beetle, Tribolium castaneum. J. Chem. Ecol. 37, 525– 532. 10.1007/s10886-011-9942-3

Đukić, N., Andrić, G., Glinwood, R., Ninkovic, V., Andjelković, B., Radonjić, A., 2021. The effect of 1-pentadecene on *Tribolium castaneum* behaviour: Repellent or attractant? Pest Manag. Sci. 77, 4034–4039. 10.1002/ps.6428

Đukić, N., Andrić, G., Ninkovic, V., Pražić Golić, M., Kljajić, P., Radonjić, A., 2020. Behavioural responses of *Tribolium castaneum* (Herbst) to different types of uninfested and infested feed. Bull. Entomol. Res. 110, 550–557. 10.1017/S0007485320000024

El-Desouky, T.A., Elbadawy, S.S., Hussain, H.B.H., Hassan, N.A., 2018. Impact of Insect Densities Tribolium Castaneum on the Benzoquinone Secretions and Aflatoxins Levels in Wheat Flour During Storage Periods. Open Biotechnol. J. 12, 104–111. 10.2174/1874070701812010104

El-Mofty, M.M., Khudoley, V.V., Sakr, S.A., Fathala, N.G., 1992. Flour infested with Tribolium castaneum, biscuits made of this flour, and 1,4-benzoquinone induce neoplastic lesions in Swiss albino mice. Nutr. Cancer 17, 97–104. 10.1080/01635589209514176

Fardet, A., 2010. New hypotheses for the health-protective mechanisms of whole-grain cereals: what is beyond fibre? Nutr. Res. Rev. 23, 65–134. 10.1017/S0954422410000041

Gao, F., Zhang, Q., Hamadou, A.H., Xu, B., 2024. The feeding preference of red flour beetle *Tribolium castaneum* Herbst (Coleoptera: Tenebrionidae) on wheat flour stored at varied temperatures: The perspective of volatile components. J. Stored Prod. Res. 108, 102397. 10.1016/j.jspr.2024.102397

Geng, P., Harnly, J.M., Chen, P., 2015. Differentiation of Whole Grain from Refined Wheat (T. aestivum) Flour Using Lipid Profile of Wheat Bran, Germ, and Endosperm with UHPLC-HRAM Mass Spectrometry. J. Agric. Food Chem. 63, 6189–6211. 10.1021/acs.jafc.5b01599

Han, S., He, K., An, J., Qiao, M., Ke, R., Wang, X., Xu, Y., Tang, X., 2023. Detection of Specific Volatile Organic Compounds in Tribolium castaneum (Herbst) by Solid-Phase Microextraction and Gas Chromatography-Mass Spectrometry. Foods 12, 2484. 10.3390/foods12132484

Kim, J., Matsuyama, S., Suzuki, T., 2005. 4,8-Dimethyldecanal, the Aggregation Pheromone of Tribolium castaneum, is Biosynthesized Through the Fatty Acid Pathway. J. Chem. Ecol. 31, 1381–1400. 10.1007/s10886-005-5292-3

Lis, Ł.B., Bakuła, T., Baranowski, M., Czarnewicz, A., 2011. The carcinogenic effects of benzoquinones produced by the flour beetle. Pol. J. Vet. Sci. 14, 159–164. 10.2478/v10181-011-0025-8

Łukowski, A., Jagiełło, R., Robakowski, P., Adamczyk, D., Karolewski, P., 2022. Adaptation of a simple method to determine the total terpenoid content in needles of coniferous trees. Plant Sci. 314, 111090. 10.1016/j.plantsci.2021.111090

Ming, Q.-L., Lewis, S.M., 2010. Pheromone production by male Tribolium castaneum (Coleoptera: Tenebrionidae) is influenced by diet quality. J. Econ. Entomol. 103, 1915–1919. 10.1603/ec10110

Negi, A., Anandharaj, A., Kalakandan, S., Rajamani, M., 2021. A Molecular Approach for the Detection and Quantification of Tribolium castaneum (Herbst) Infestation in Stored Wheat Flour. Food Technol. Biotechnol. 59, 112–121. 10.17113/ftb.59.01.21.6902

Ney, P., Boland, W., 1987. Biosynthesis of 1-alkenes in higher plants. Eur. J. Biochem. 162, 203–211. 10.1111/j.1432-1033.1987.tb10562.x

Niu, Y., Hardy, G., Agarwal, M., Hua, L., Ren, Y., 2016. Characterization of volatiles *Tribolium castaneum* (H.) in flour using solid phase microextraction–gas chromatography mass spectrometry (SPME–GCMS). Food Sci. Hum. Wellness 5, 24– 29. 10.1016/j.fshw.2015.11.002

Pappas, P.W., Morrison, S.E., 1995. Benzoquinones of the beetles, Tribolium castaneum and Tribolium confusum. Prep. Biochem. 25, 155–168. 10.1080/10826069508010117

Phillips, T.W., 1997. Semiochemicals of stored-product insects: Research and applications. J. Stored Prod. Res., Ecologially safe alternative for the control of stored-product insects 33, 17–30. 10.1016/S0022-474X(96)00039-2

Poddar-Sarkar, M., 1996. The fixative lipid of tiger pheromone. J. Lipid Mediat. Cell Signal. 15, 89–101. 10.1016/s0929-7855(96)00547-0

Pointer, M.D., Gage, M.J.G., Spurgin, L.G., 2021. Tribolium beetles as a model system in evolution and ecology. Heredity 126, 869–883. 10.1038/s41437-021-00420-1

Pomeranz, Y., 1988. Composition and functionality of wheat flour components, in: In Wheat: Chemistry and Technology. American Association of Cereal Chemists, St. Paul, MN, pp. 219–370.

Rath, A., Benita, M., Doron, J., Scharf, I., Gottlieb, D., 2021. Social communication activates the circadian gene Tctimeless in Tribolium castaneum. Sci. Rep. 11, 16152. 10.1038/s41598-021-95588-1

Renou, M., Anton, S., 2020. Insect olfactory communication in a complex and changing world. Curr. Opin. Insect Sci., Neuroscience * Biomechanics of Insect Flight and Bio-inspired engineering 42, 1–7. 10.1016/j.cois.2020.04.004

Seifelnasr, Y.E., Hopkins, T.L., Mills, R.B., 1982. Olfactory responses of adultTribolium castaneum (Herbst), to volatiles of wheat and millet kernels, milled fractions, and extracts. J. Chem. Ecol. 8, 1463–1472. 10.1007/BF00989103

Semeao, A.A., Campbell, J.F., Whitworth, R.J., Sloderbeck, P.E., 2011. Response of *Tribolium castaneum* and *Tribolium confusum* adults to vertical black shapes and its potential to improve trap capture. J. Stored Prod. Res. 47, 88–94. 10.1016/j.jspr.2011.01.002

Shamjana, U., Grace, T., Shamjana, U., Grace, T., 2021. Review of Insecticide Resistance and Its Underlying Mechanisms in <em>Tribolium castaneum</em>. IntechOpen. 10.5772/intechopen.100050

Stevenson, B.J., Cai, L., Faucher, C., Michie, M., Berna, A., Ren, Y., Anderson, A., Chyb, S., Xu, W., 2017. Walking Responses of Tribolium castaneum (Coleoptera: Tenebrionidae) to Its Aggregation Pheromone and Odors of Wheat Infestations. J. Econ. Entomol. 110, 1351–1358. 10.1093/jee/tox051

Suh, E., Bohbot, J.D., Zwiebel, L.J., 2014. Peripheral olfactory signaling in insects. Curr. Opin. Insect Sci. 6, 86–92. 10.1016/j.cois.2014.10.006

Suzuki, T., 1980. 4,8-Dimethyldecanal: The Aggregation Pheromone of the Flour Beetles, *Tribolium castaneum* and *T. confusum* (Coleoptera: Tenebrionidae). Agric. Biol. Chem. 44, 2519–2520. 10.1080/00021369.1980.10864359

Tena, N., Lobo-Prieto, A., Aparicio, R., García-González, D.L., 2019. Storage and Preservation of Fats and Oils, in: Ferranti, P., Berry, E.M., Anderson, J.R. (Eds.), Encyclopedia of Food Security and Sustainability. Elsevier, Oxford, pp. 605–618. 10.1016/B978-0-08-100596-5.22268-3

Tian, X., Hao, J., Wu, F., Hu, H., Zhou, G., Liu, X., Zhang, T., 2022. 1-Pentadecene, a volatile biomarker for the detection of *Tribolium castaneum* (Herbst) (Coleoptera: Tenebrionidae) infested brown rice under different temperatures. J. Stored Prod. Res. 97, 101981. 10.1016/j.jspr.2022.101981

Trebels, B., Dippel, S., Goetz, B., Graebner, M., Hofmann, C., Hofmann, F., Schmid, F.-R., Uhl, M., Vuong, M.-P., Weber, V., Schachtner, J., 2021. Metamorphic development of the olfactory system in the red flour beetle (Tribolium castaneum, Herbst). BMC Biol. 19, 155. 10.1186/s12915-021-01055-8

Villaverde, M.L., Juárez, M.P., Mijailovsky, S., 2007. Detection of *Tribolium castaneum* (Herbst) volatile defensive secretions by solid phase microextraction–capillary gas chromatography (SPME-CGC). J. Stored Prod. Res. 43, 540–545. 10.1016/j.jspr.2007.03.003

Wagner, K., Aikins, M.J., Phillips, T., 2018. Preferences of the Red Flour Beetle, Tribolium castaneum, for Nutritionally Different Dog Foods.

Waldbauer, G.P., Bhattacharya, A.K., 1973. Self-selection of an optimum diet from a mixture of wheat fractions by the larvae of *Tribolium confusum*. J. Insect Physiol. 19, 407– 418. 10.1016/0022-1910(73)90115-7

Wang, V.-S., 1981. The isolation and identification of volatile compounds in wheat which induce an aggregation response to the red flour beetle, Tribolium casteneum (Herbst.). Kansas State University.

Xi, Y., Ikram, S., Zhao, T., Shao, Y., Liu, R., Song, F., Sun, B., Ai, N., 2023. 2-Heptanone, 2-nonanone, and 2-undecanone confer oxidation off-flavor in cow milk storage. J. Dairy Sci. 106, 8538–8550. 10.3168/jds.2022-23056

Xu, X., Zhou, L., Yu, H., Sun, G., Fei, S., Zhu, J., Ma, Y., 2024. Winter wheat ear counting based on improved YOLOv7x and Kalman filter tracking algorithm with video streaming. Front. Plant Sci. 15. 10.3389/fpls.2024.1346182

Yemm, E.W., Willis, A.J., 1954. The estimation of carbohydrates in plant extracts by anthrone. Biochem. J. 57, 508–514. 10.1042/bj0570508

