## Supplemental for "Unveiling the chemical and behavioural ecology of *Tribolium castaneum* (Herbst, 1797) in wheat flour: Alterations in flour metabolic content and the role of chemical cues in modulating beetles’ behaviour and regulating population growth"

***Supplementary file***

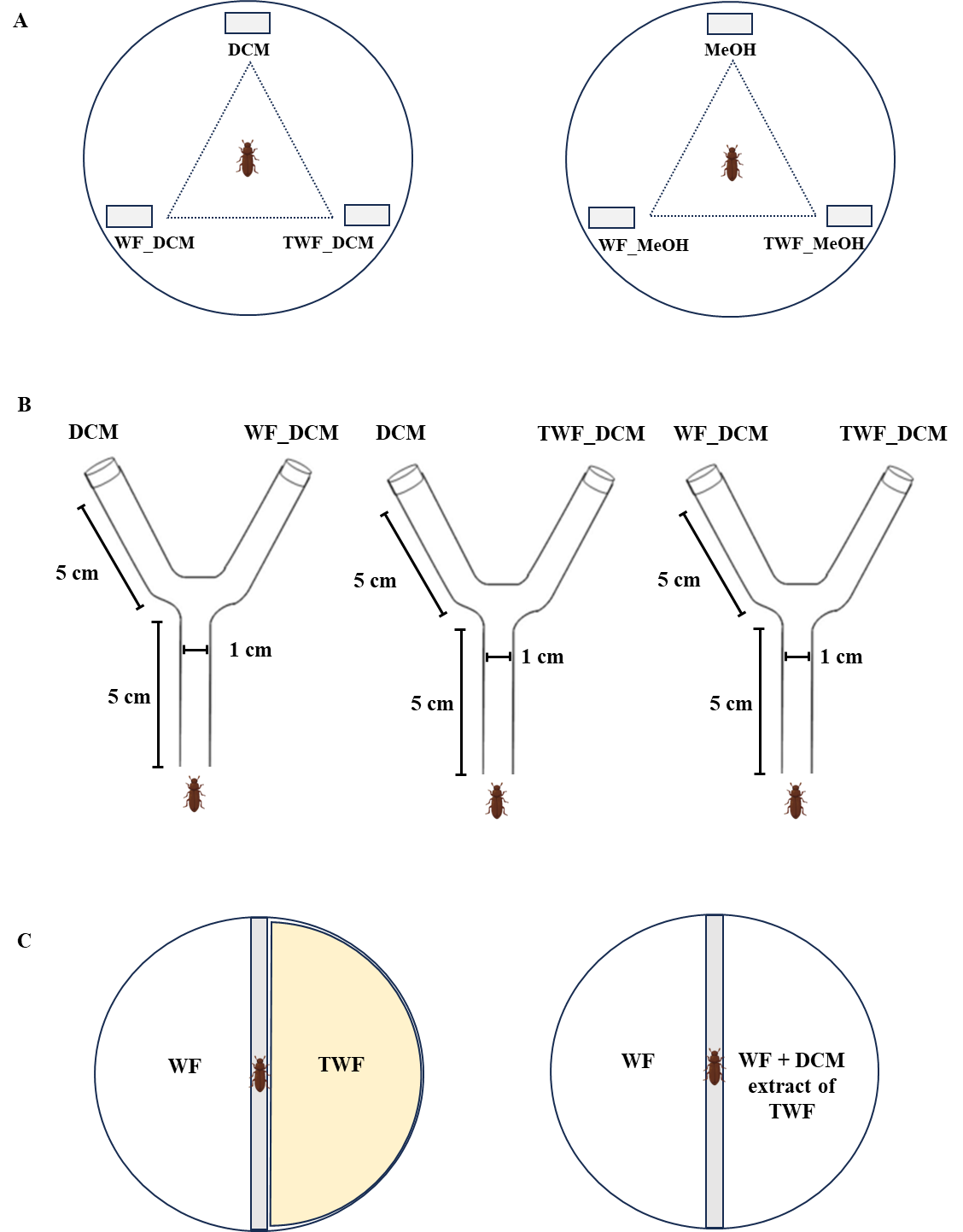

**Supplementary Fig.1.** Experimental setups to analyse behavioural modulation activity of WF and TWF extract on *T. castaneum*.

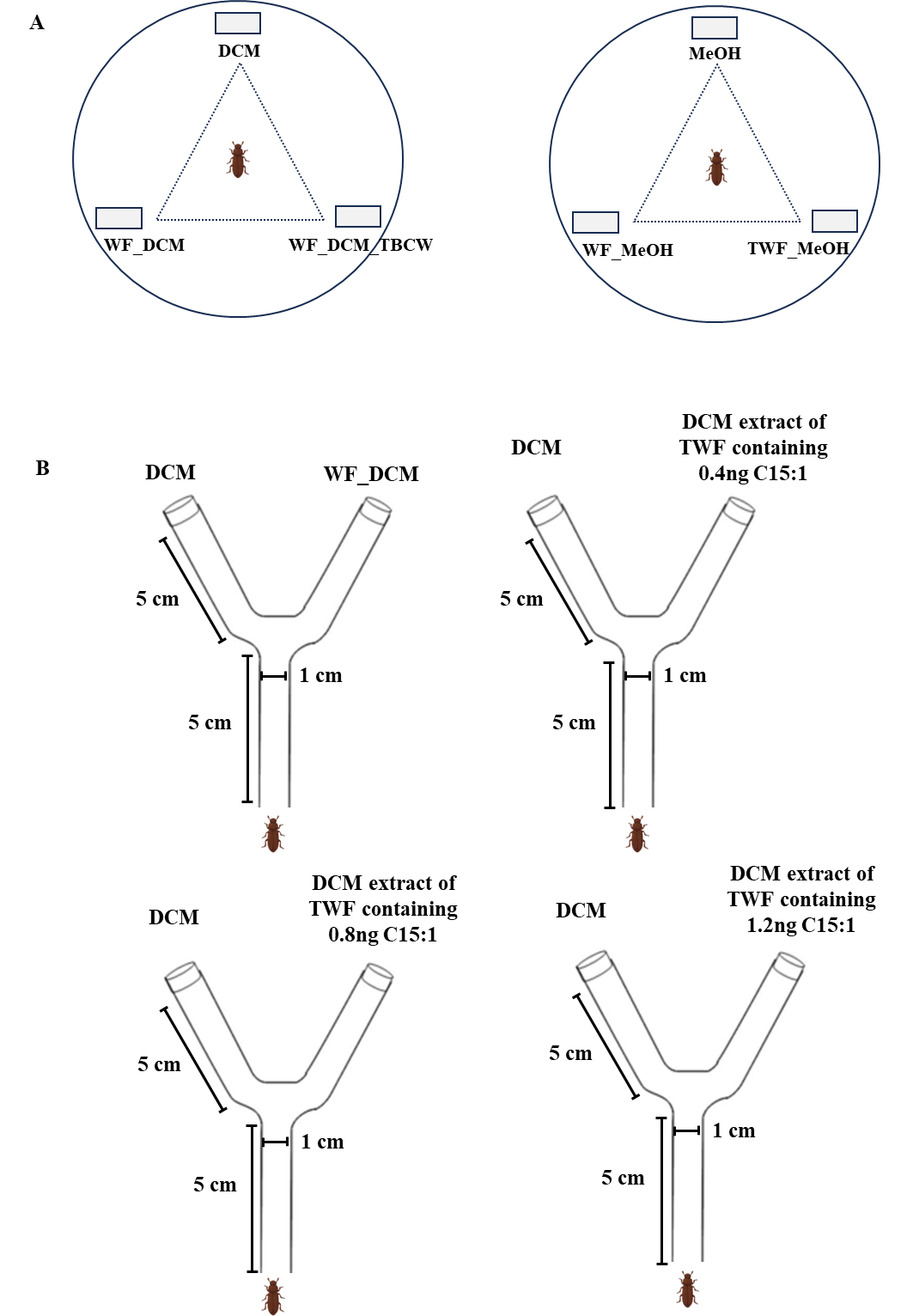

**Supplementary Fig.2.** Experimental setups to analyse the role of infested flour odour with different concentrations of 1-Pentadecene on *T. castaneum* preference behaviour.

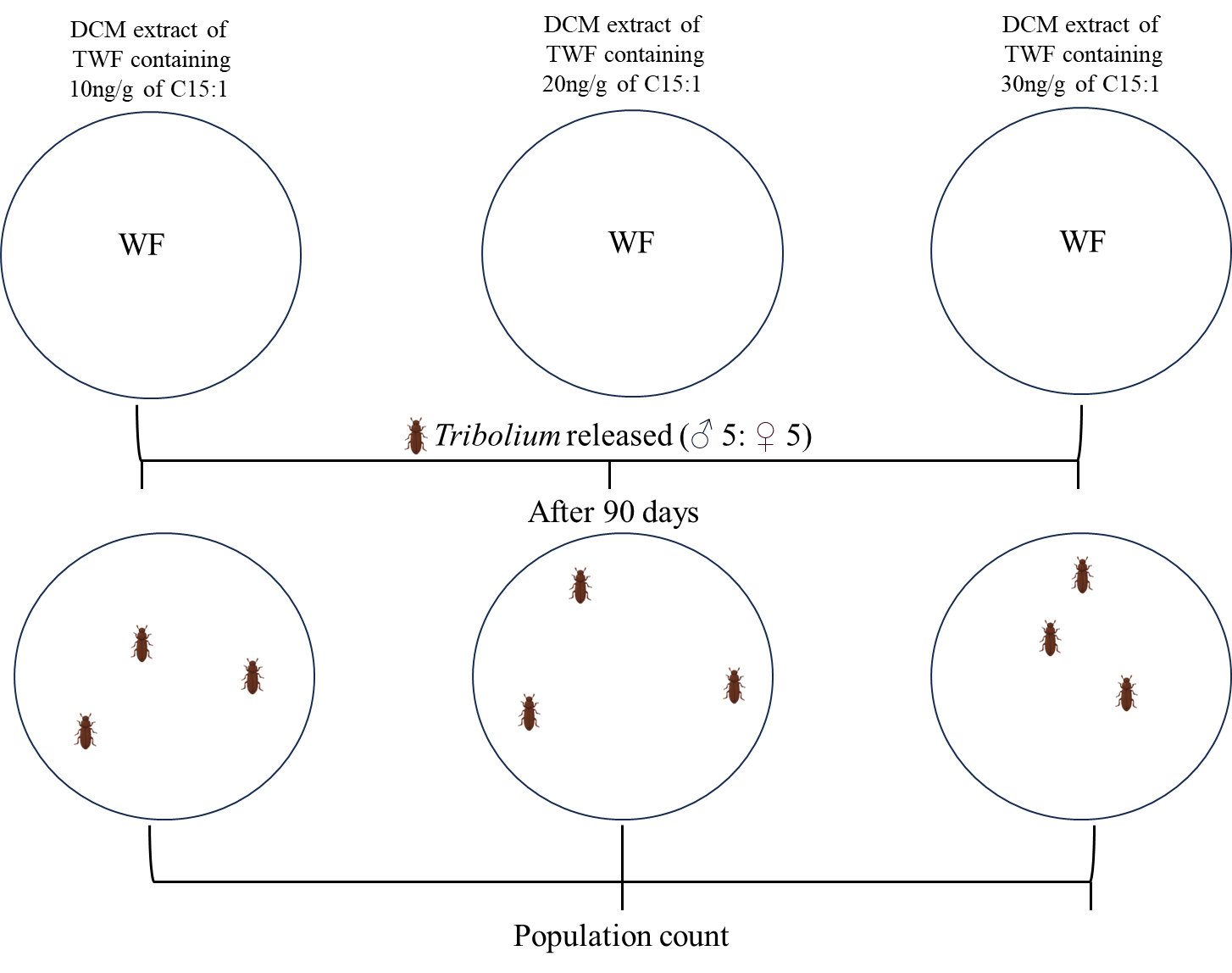

**Supplementary Fig.3.** Experimental setups to analyse the effects of *T. castaneum* odorants in determining reproductive success.

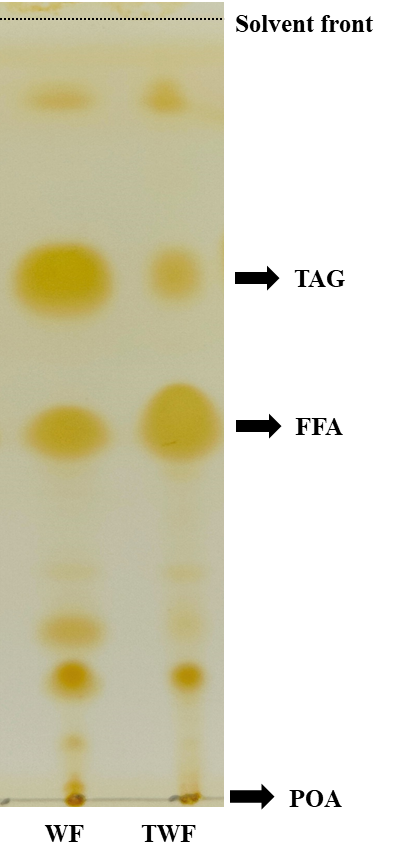

**Supplementary Fig. 4.** Thin layer chromatography (TLC) for analysis of Neutral lipid extracted from total lipid of WF and TWF developed in Petroleum ether : diethyl ether : acetic acid (80:20:2) and stained with Iodine vapour. FFA: Free Fatty Acid; TAG: Triacylglycerol; POA – point of application (R_f_ of FFA- 0.47; R_f_ of TAG- 0.66). [Thin Layer Chromatographic plates- E.Merck, Germany; Silica gel ­60F_254_; layer thickness 200 µm]

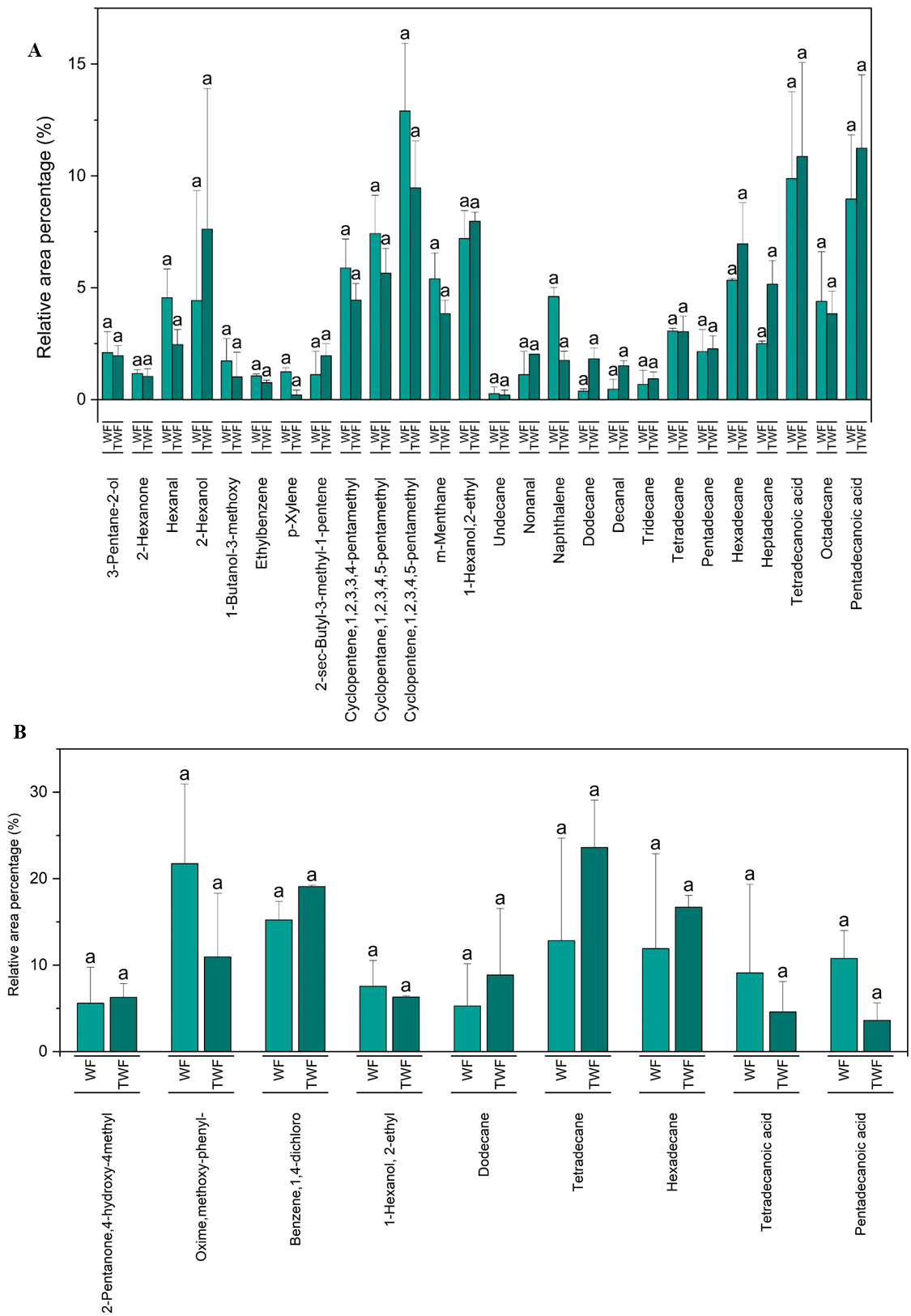

**Supplementary Fig. 5.** The bar graphs illustrate the relative abundance of commonly detected VOCs in the DCM extract (A) and MeOH extract (B) from WF and TWF. Letters above the bar plots indicate statistical difference, where plots sharing the same letter show no significant difference.

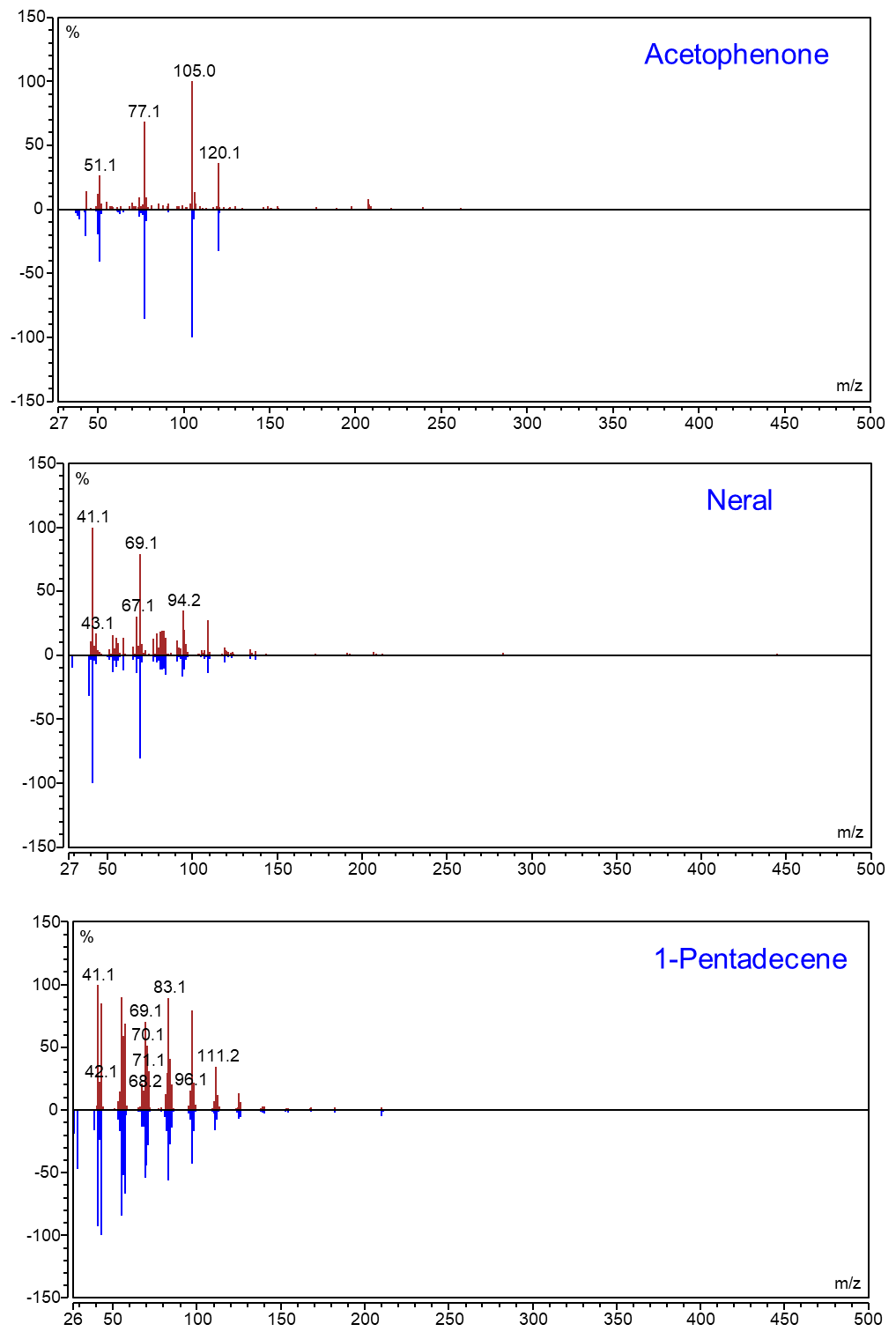

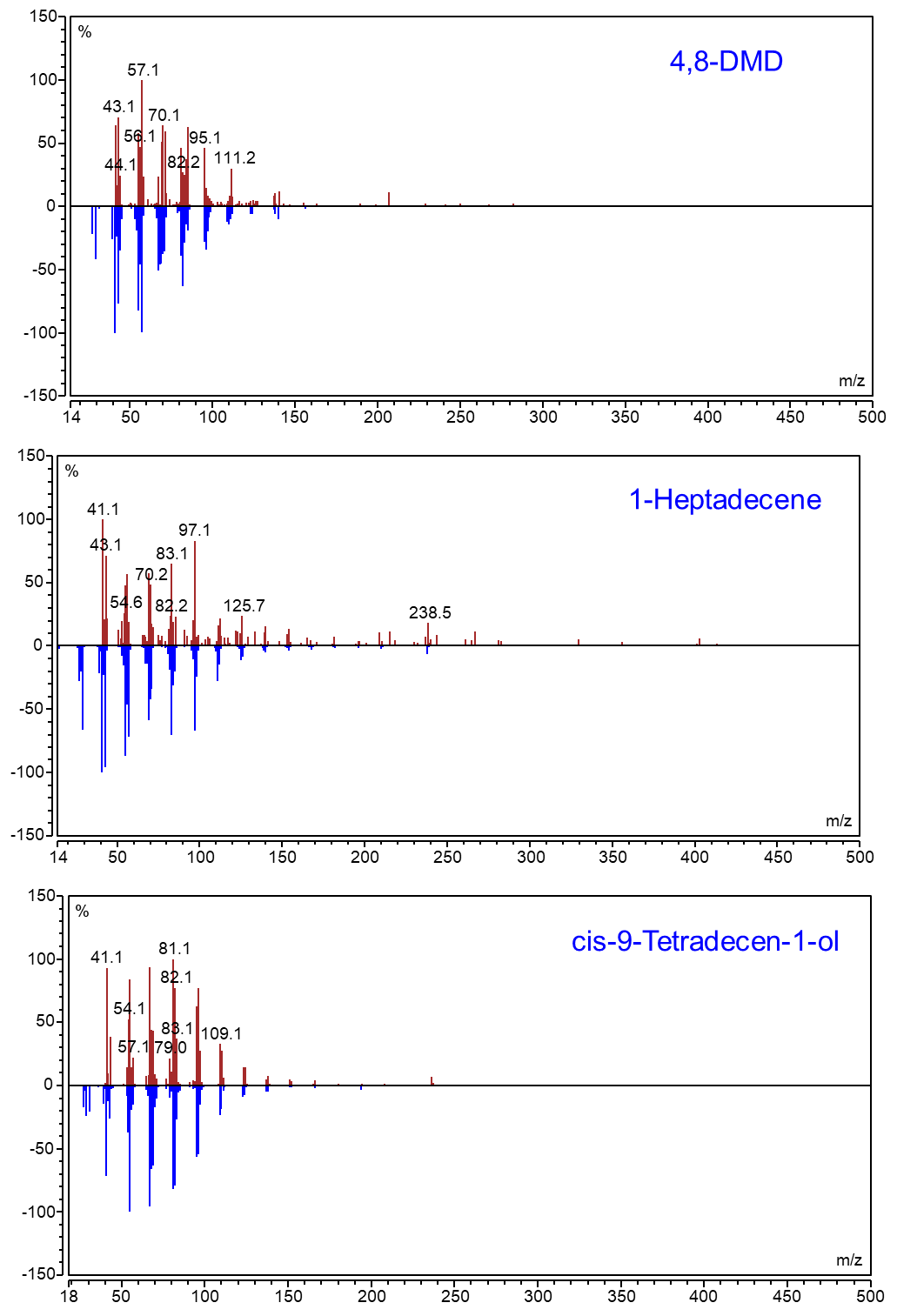

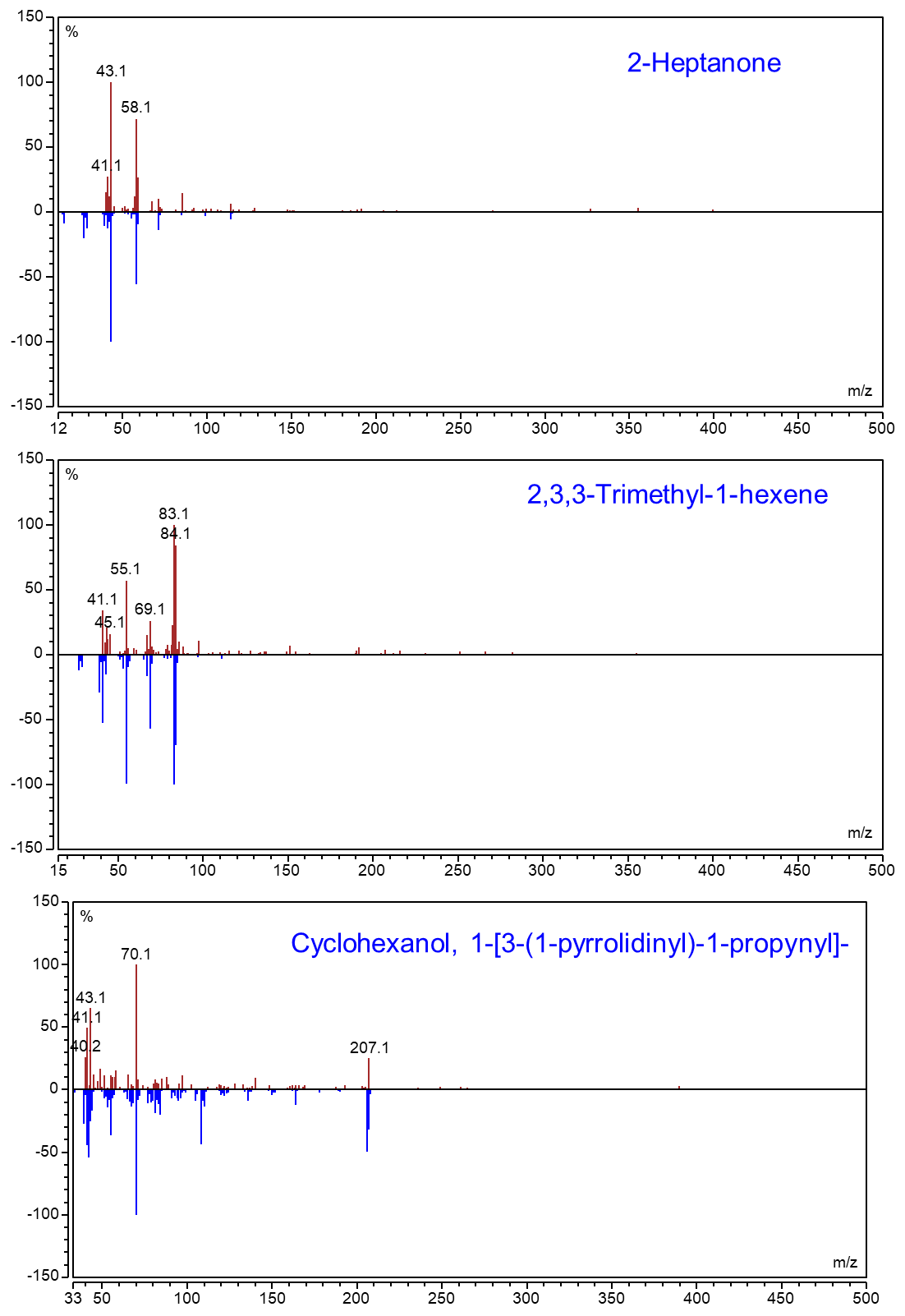

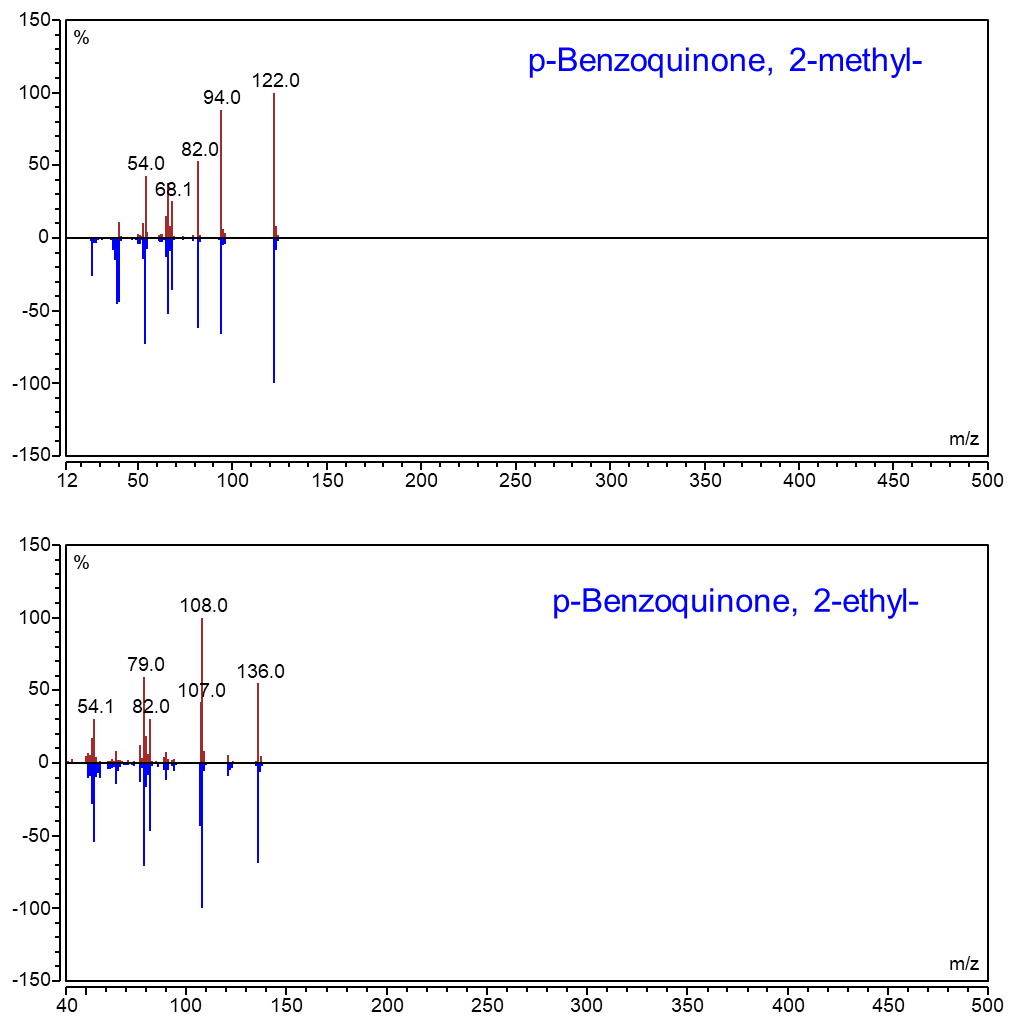

**Supplementary Fig. 6. Mass fragmentation patterns of some important odorants (red – sample compound mass; blue- NIST database)**

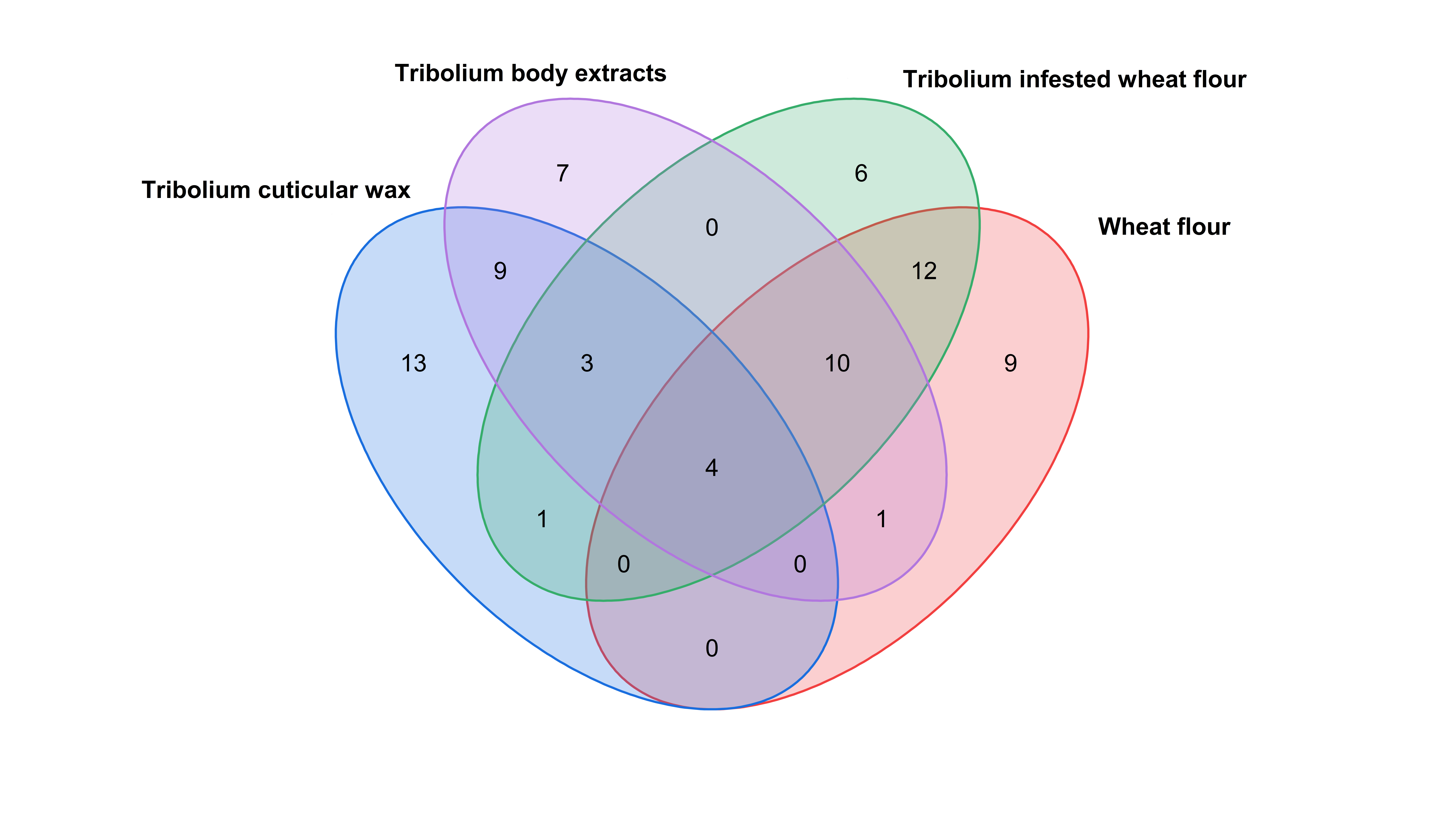

**Supplementary Fig. 7.** This Venn diagram illustrates the overlapping compounds found in *Tribolium* body extracts, *Tribolium-*infested wheat flour, *Tribolium* cuticular wax, and fresh wheat flour. The numbers in each region represent the number of compounds unique or shared between different sample types

17 compounds are common to Tribolium body extracts and Tribolium infested wheat flour. 8 compounds are common to Tribolium cuticular wax, and Tribolium infested wheat flour. 7 compounds are common to Tribolium body extracts, Tribolium cuticular wax, and Tribolium infested wheat flour.

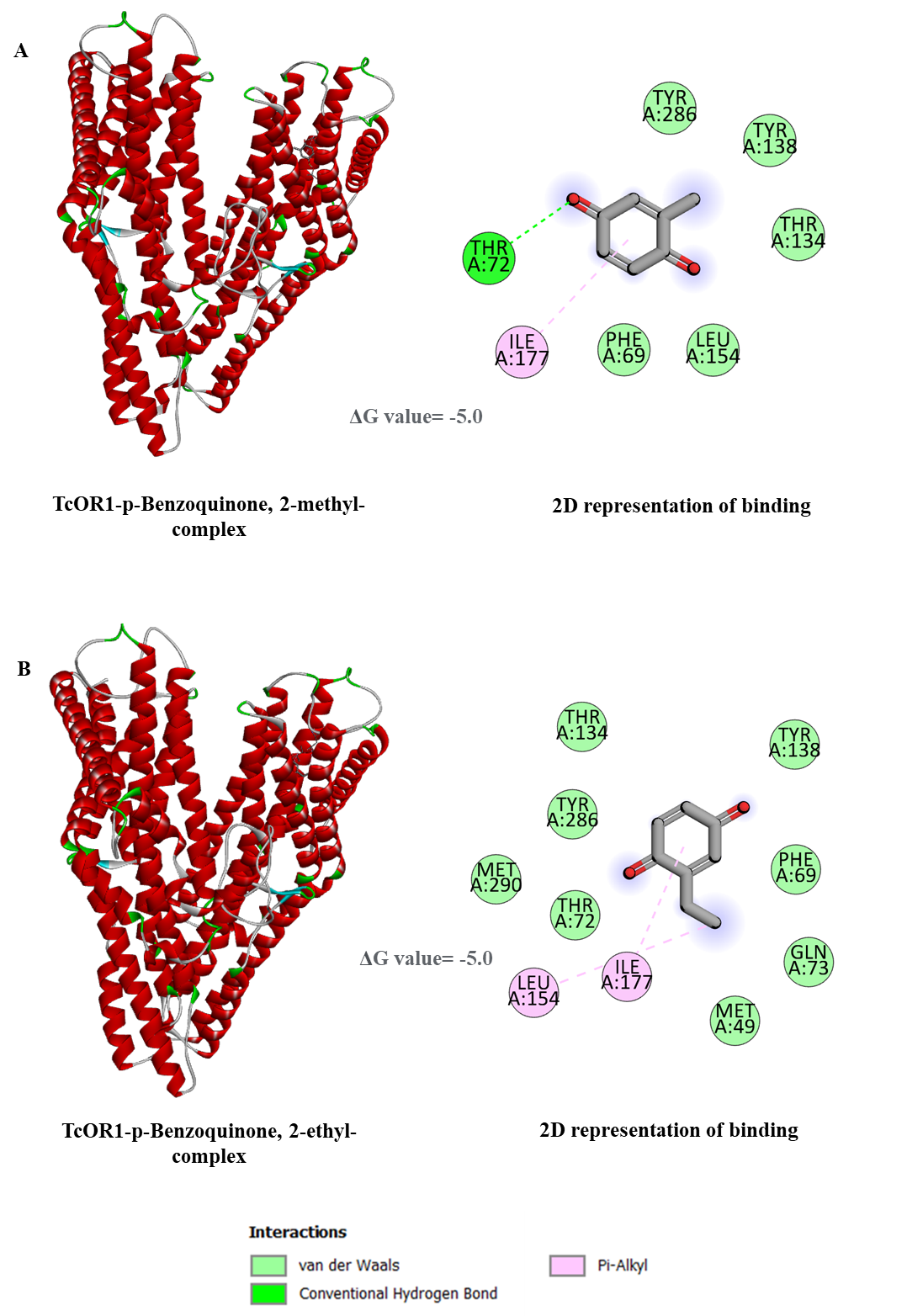

**Supplementary Fig. 8.** *In-silico* molecular docking visualization of TcOR1 (odorant receptor) in complex with p-Benzoquinone-2-methyl (A) and p-Benzoquinone-2-ethyl (B), using Discovery Studio Visualizer v21.1.0.20298. The 3D protein-ligand interaction maps are shown with the receptor in red helical ribbons. Ligand-binding details are displayed in close-up insets, along with 2D diagrams highlighting key residues involved in interactions. ΔG values are mentioned in the diagram.

**Supplementary Table 1.** Amount of 4,8-DMD and 1-Pentadecene in TWF in different experimental sets.

|  | Amount in TWF (ng/g) | |
| --- | --- | --- |
|  | 4,8-DMD | 1-Pentadecene |
| Experiment set 1 | 6.90±0.75 | 27.52±7.56 |
| Experiment set 2 | 7.38±1.69 | 19.06±6.64 |
| Experiment set 3 | 6.16±2.28 | 10±6.47 |

**Supplementary Table 2.** List of fatty acids identified from WF and TWF [amount in relative percentage].

|  | Relative abundance (%) [Mean±SD] | |
| --- | --- | --- |
| Fatty Acids | WF | TWF |
| C12:0 | 0.03 ± 0.010 | 0.16±0.042 |
| C14:0 | 0.15 ±0.111 | 0.34±0.013 |
| C15:0 | 0.20 ± 0.142 | 0.33±0.023 |
| C16:0 | 36.82±2.141 | 49.73±0.197 |
| C17:0 | 0.22±0.102 | 0.25±0.022 |
| C18:0 | 2.02±0.535 | 3.14±0.037 |
| C20:0 | 0.50±0.032 | 0.24±0.090 |
| C21:0 | 0.30±0.223 | 0.00±0.000 |
| C22:0 | 0.36±0.065 | 0.26±0.074 |
| C23:0 | 0.14±0.002 | 0.00±0.000 |
| C24:0 | 0.34±0.025 | 0.00 |
| **⅀SFA** | 41.08±3.389 | 54.44±0.499 |
| C16:1 | 0.16±0.141 | 0.10±0.040 |
| C17:1 | 0.09±0.011 | 0.00±0.000 |
| C18:2 | 36.50±3.485 | 23.43±0.408 |
| C18:1(11) | 12.47±0.509 | 9.79±0.036 |
| C18:1(9) | 7.20±1.801 | 10.94±0.151 |
| C20:1 | 0.79±0.438 | 1.29±0.031 |
| C22:1 | 0.05±0.011 | 0.00 |
| **⅀UFA** | 57.26±6.396 | 45.56±0.667 |

⅀SFA = Summation of saturated fatty acids; ⅀UFA= Summation of unsaturated fatty acids.

**Supplementary Table 3.** List of identified head space volatiles (HSVs) of WF and TWF obtained from HS-SPME technique from WF and TWF.

| Rt (Min) | Compounds | WF (HS-SPME) | TWF (HS-SPME) | LRI |
| --- | --- | --- | --- | --- |
| 3.302 | 1-Pentanol | ✓ | ✓ | 757 |
| 3.758 | Hexanal | ✓ | ✓ | 797 |
| 4.546 | 2-Heptanone |  | ✓ | 843 |
| 4.845 | Ethylbenzene | ✓ | ✓ | 861 |
| 4.935 | 1-Hexanol | ✓ | ✓ | 866 |
| 4.996 | p-Xylene | ✓ | ✓ | 869 |
| 5.402 | Styrene | ✓ | ✓ | 893 |
| 5.529 | Oxime, methoxy-phenyl | ✓ | ✓ | 900 |
| 5.821 | Butyrolactone | ✓ | ✓ | 914 |
| 6.864 | Benzaldehyde | ✓ | ✓ | 963 |
| 7.202 | Hexanoic acid |  | ✓ | 979 |
| 7.394 | 6-Methyl-5-hepten-2-one |  | ✓ | 988 |
| 7.511 | Furan, 2-pentyl | ✓ | ✓ | 993 |
| 7.679 | Decane |  | ✓ | 1001 |
| 7.736 | Octanal | ✓ | ✓ | 1003 |
| 8.011 | Benzene,1,4-dichloro | ✓ | ✓ | 1016 |
| 8.329 | 1-Hexanol,2-ethyl |  | ✓ | 1030 |
| 9.952 | Undecane | ✓ | ✓ | 1101 |
| 10.05 | Nonanal | ✓ | ✓ | 1105 |
| 10.365 | Hexanoic acid,2-ethyl |  | ✓ | 1119 |
| 11.518 | Benzoic acid | ✓ | ✓ | 1171 |
| 12.196 | Dodecane | ✓ | ✓ | 1201 |
| 14.345 | Tridecane | ✓ | ✓ | 1300 |
| 15.077 | **4,8-Dimethyldecanal** |  | ✓ | 1336 |
| 15.221 | Benzamide | ✓ |  | 1343 |
| 16.391 | Tetradecane | ✓ | ✓ | 1400 |
| 18.182 | **1-Pentadecene** |  | ✓ | 1493 |

Several compounds are commonly detected in solvent runs and SPME, including two key compounds: 4,8-Dimethyldecanal and 1-Pentadecene (bold).

**Supplementary Table 4.** PC loading values of all VOCs identified from WF and TWF. Top ten compounds with high loading values are demarcated in colour.

| **Compounds** | **PC 1** | **PC 2** | **PC 3** | **PC 4** | **PC 5** |
| --- | --- | --- | --- | --- | --- |
| 1-Pentadecene | 0.672 | 0.19256 | -0.016 | 0.53256 | -0.0975 |
| cis-9-Tetradecen-1-ol | 0.46884 | 0.12511 | 0.05606 | -0.2191 | 0.20179 |
| 1-Heptadecene | 0.11436 | 0.02936 | 0.0221 | 0.06127 | -0.0098 |
| 2-Hexanol | 0.1088 | -0.4317 | -0.8061 | 0.02762 | 0.11434 |
| 4,8-Dimethyldecanal | 0.09646 | 0.03158 | -0.031 | 0.03478 | 0.01529 |
| Nonadecane | 0.08963 | 0.02872 | -0.0242 | 0.0031 | 0.02717 |
| Heptadecane | 0.08264 | 0.01921 | 0.10508 | -0.2002 | 0.29659 |
| Neral | 0.07955 | 0.02324 | -0.0051 | -0.0459 | 0.04302 |
| Dodecane | 0.05313 | 0.01096 | -0.0599 | 0.01365 | -0.0453 |
| 2-Heptanone | 0.04588 | 0.01702 | -0.0293 | 0.02405 | 0.00804 |
| 1-Butanol-3-methoxy | 0.04241 | -0.0039 | 0.11573 | -0.0561 | 0.00217 |
| Pentadecanoic acid | 0.03509 | -0.3713 | 0.27681 | 0.18209 | 0.4301 |
| Decanal | 0.03298 | 0.06177 | -0.0207 | -0.0831 | 0.17531 |
| 2,3,3-Trimethyl-1-hexene | 0.03279 | 0.01124 | -0.0142 | 0.04926 | -0.0121 |
| Hexadecane | 0.01763 | -0.0182 | 0.19071 | -0.0987 | -0.0033 |
| Nonanal | 0.01751 | 0.12135 | 0.01764 | 0.03314 | 0.07199 |
| 2-sec-Butyl-3-methyl-1-pentene | 0.01401 | 0.12806 | -0.0385 | 0.05908 | 0.0756 |
| Acetophenone | 0.01188 | -0.0011 | 0.0324 | 0.05082 | -0.0321 |
| Cyclohexanol,1-[3-(1-pyrrolidinyl)-1-propynyl] | 0.00283 | -0.0003 | 0.00772 | -0.0949 | 0.04494 |
| Tridecane | -0.0006 | 0.06585 | 0.04583 | -0.0178 | 0.06282 |
| Undecane | -0.0022 | -0.0373 | 0.01749 | 0.03947 | -0.1249 |
| Tetradecanoic acid | -0.0147 | -0.5157 | 0.35963 | 0.36602 | -0.1653 |
| 2-Hexanone | -0.0179 | 0.01829 | -0.0335 | -0.061 | -0.1808 |
| Ethylbenzene | -0.0209 | -0.0168 | -0.0134 | 0.14099 | 0.04907 |
| Pentadecane | -0.0215 | 0.09322 | 0.08313 | -0.0318 | -0.2726 |
| 3-Hexen-1-ol, (Z) | -0.022 | -0.0214 | -0.0013 | -0.0146 | -0.1242 |
| 1-Hexanol, 2-ethyl | -0.0278 | -0.1503 | -0.08 | 0.11177 | -0.2224 |
| Hexanoic acid | -0.0291 | -0.0999 | -0.0155 | -0.032 | -0.1613 |
| Tetradecane | -0.0292 | -0.0078 | 0.07569 | -0.1333 | -0.0758 |
| 3-Penten-2ol | -0.033 | 0.10016 | -0.0287 | -0.0209 | 0.11613 |
| Heptane | -0.0403 | -0.1383 | -0.0214 | 0.02264 | 0.1456 |
| Octadecane | -0.0436 | -0.2822 | 0.05915 | -0.1719 | -0.0867 |
| p-Xylene | -0.0533 | 0.00094 | 0.03122 | 0.08714 | -0.0528 |
| Acetic acid, 2-ethylhexyl ester | -0.0676 | -0.0452 | -0.0002 | 0.05663 | 0.15737 |
| Nonanoic acid | -0.0715 | 0.02529 | 0.0138 | 0.02885 | -0.0788 |
| Vanillin | -0.0807 | 0.0314 | 0.01613 | 0.09507 | 0.25234 |
| 1-Butanol,3-methoxy | -0.0919 | 0.08244 | 0.0273 | 0.0677 | 0.01672 |
| 3-Hydroxy-3-methyl-2-butanone | -0.0983 | 0.02255 | 0.01664 | 0.0806 | 0.12952 |
| m-Menthane | -0.1214 | 0.09844 | -0.0366 | 0.24297 | -0.1071 |
| Cyclopentene,1,2,3,3,4-pentamethyl | -0.1226 | 0.11536 | -0.0509 | -0.0396 | -0.186 |
| 1,1-Dimethyl-3-chloropropanol | -0.1228 | 0.03304 | 0.02172 | 0.10378 | 0.17382 |
| Hexanal | -0.1358 | 0.11005 | -0.0393 | 0.1206 | 0.06906 |
| Carbonic acid, ethyl phenyl ester | -0.1535 | 0.11184 | 0.04065 | 0.14121 | 0.20906 |
| Cyclopentane,1,2,3,4,5-pentamethyl | -0.154 | 0.15625 | -0.081 | 0.22899 | -0.1602 |
| Naphthalene | -0.1591 | -0.001 | -0.028 | 0.27584 | 0.14491 |
| Cyclopentene,1,2,3,4,5-pentamethyl | -0.283 | 0.27691 | -0.1578 | 0.24405 | 0.18115 |

**Supplementary Table 5.** Grid box values inside TcOR1 during volatile-OR docking

| center_x | -5.65225604853 |
| --- | --- |
| center_y | 7.89493257979 |
| center_z | -25.821024394 |
| size_x | 23.0570797778 |
| size_y | 17.8895331534 |
| size_z | 26.2426386375 |

**Supplementary Table 6.** Ten distinct compounds detected in TWF are found in various insects, functioning as semiochemicals. The table provides the molecular weight of each compound, and their presence in different insect groups. The listed references support the presence and role of these compounds in specific insect species, as documented by multiple researchers.

| Name of Compounds | Presence in Order Coleoptera | Presence in other orders | References |
| --- | --- | --- | --- |
| Acetophenone (M.W. =120.15 ) | *Dendroctonus ponderosae* ^P,A^ ,  *Dendroctonus pseudotsugae* ^R,A^ , *Dendroctonus rufipennis* ^A^ , *Dryocoetes confuses* ^A^*, Taphrorychus bicolor* ^P^ | *Cerapachys jacobsoni* ^P^ (Hymenoptera)*, Vespa velutina* ^P^ (Hymenoptera) | (Francke et al., 1996; Gries et al., 1992; Kohnle et al., 1987; Morgan et al., 2008; Pureswaran et al., 2004; Pureswaran and Borden, 2004; Thiéry et al., 2018) |
| Neral  (M.W. =152.24) | *Bledius dissimilis* ^R^*,*  *B. furcatus* ^R^ *,*  *B. mandibularis* ^R^, *Megacyllene caryae* ^P^, *Platypus koryoensis* ^P^ | *Andrena denticulata*^P^ (Hymenoptera)*, Centris adani* ^P^ (Hymenoptera),  *Pieris napi* ^p^ (Lepidoptera) | (Kim et al., 2009; Lacey et al., 2008; Steidle and Dettner, 1995; Tengö and Bergstrom, 1977; Vinson et al., 1982; Wheeler et al., 1972) |
| 4,8-DMD  (M.W.= 184.32) | *Tribolium audax* ^P^*,*  *T. brevicornis* ^P^*,*  *T. castaneum* ^P^*,*  *T. confusum* ^P^*,*  *T. destructor* ^P^*,*  *T. ferrugineum* ^P^*,*  *T. freeman* ^P^*,*  *T. madens* ^P^ |  | (Arnaud et al., 2002) |
| 1-Pentadecene  (M.W.= 210.4) | *Bembidion interventor* ^A^, *T. audax* ^P^*, T. brevicornis* ^P^*, T. castaneum* ^P^*,*  *T. confusum* ^P^*,*  *T. destructor* ^P^*,*  *T. ferrugineum* ^P^*,*  *T. freeman* ^P^*, T. madens* ^P^ | *Nannotrigona testaceicornis* ^P^ (Hymenoptera), *Solenopsis geminate* ^P^ (Hymenoptera) | (Arnaud et al., 2002; Cruz-López et al., 2001; Evans, 1988; Pianaro et al., 2009) |
| 1-Heptadecene (M.W.=238.46) | *Oxelytrum discicolle* ^P^, *Cibdelis gibbose* ^AL^, *Bembidion interventor* ^A^, *Cratidus osculans* ^AL^ | *Nothomyrmecia macrops* ^AL^ (Hymenoptera), *Formica polyctena* ^P^ (Hymenoptera) | (Billen et al., 1988; Evans, 1988; Fockink et al., 2013; Löfqvist and Bergström, 1980; Tschinkel, 1975) |
| 2-Heptanone  (MW-114.19) | *Aethina tumida* ^A^, *Ontholestes murinus* ^AL^, *Anaglyptus mysticus* ^P^, *Rhynchophorus palmarum* ^K^ | *Hypoclinea bidens* ^P^ (Hymenoptera), *Iridomyrmex pruunosus* ^AL^ (Hymenoptera) | (Blum et al., 1982; Huth and Dettner, 1990; Molander et al., 2019; Rochat et al., 2000; Scheffrahn et al., 1984; Torto et al., 2005) |
| cis-9-Tetradecen- 1-ol  (M.W.= 212.37) |  | *Psithyrus rupestris* ^P^ (Hymenoptera), *Spodoptera eridania* ^P^ (Lepidoptera), *Graphania mutans* ^P^, (Lepidoptera) | (Frérot and Foster, 1991; Lanne et al., 1987; Teal et al., 1985) |
| Nonadecane (M.W.= 268.51) | *Rhynchophorus ferrugineus*^P^*,* *Eusphalerum abdominale*^AL^*, Eusphalerum sorbi*^AL^*, Eusphalerum stramineum*^AL^ | *Myrmica scabrinodis*^P^ (Hymenoptera)*, Colletes fodiens*^P^ (Hymenoptera)*, Formica nigricans*^P^ (Hymenoptera)*, Toxotrypana curvicauda*^A^ (Diptera) | (A. and Srour, 2009; Bergström and Tengö, 1978; Dettner and Schwinger, 1977; Morgan et al., 1979) |
| 2,3,3-Trimethyl-1-hexene  (M.W.= 126.24) |  |  |  |
| Cyclohexanol,1-[3-(1-pyrrolidinyl)-1-propynyl]  (M.W= 207.31) |  |  |  |

The molecular weights (M.W.) of the compounds are indicated in the parentheses. The symbols A, R, P, K, and AL represent Attractants, Repellents, Pheromones, Kairomone, and Allomones, respectively.

**References**

A., M.M., Srour, H.A., 2009. Desiccation intolerance of the red palm weevil, Rhynchophorus ferrugineus (oliv) adults in relation to their cuticular hydrocarbons. Egyptian Academic Journal of Biological Sciences. A, Entomology 2, 47–53. https://doi.org/10.21608/eajbsa.2009.15452

Arnaud, L., Lognay, G., Verscheure, M., Leenaers, L., Gaspar, C., Haubruge, E., 2002. Is Dimethyldecanal a Common Aggregation Pheromone of Tribolium Flour Beetles? J Chem Ecol 28, 523–532. https://doi.org/10.1023/A:1014587927784

Bergström, G., Tengö, J., 1978. Linalool in mandibular gland secretion of Colletes bees (Hymenoptera: Apoidea). J Chem Ecol 4, 437–449. https://doi.org/10.1007/BF00989500

Billen, J.P.J., Jackson, B.D., Morgan, E.D., 1988. Secretion of the Dufour gland of the ant Nothomyrmecia macrops (Hymenoptera:Formicidae). Experientia 44, 715–719. https://doi.org/10.1007/BF01941041

Blum, M.S., Jones, T.H., Snelling, R.R., Overal, W.L., Fales, H.M., Highet, R.J., 1982. Systematic implications of the exocrine chemistry of some Hypoclinea species. Biochemical Systematics and Ecology 10, 91–94. https://doi.org/10.1016/0305-1978(82)90057-6

Cruz-López, L., Rojas, J.C., De La Cruz-Cordero, R., Morgan, E.D., 2001. Behavioral and Chemical Analysis of Venom Gland Secretion of Queens of the Ant Solenopsis geminata. J Chem Ecol 27, 2437–2445. https://doi.org/10.1023/A:1013671330253

Dettner, K., Schwinger, G., 1977. Microdetermination of defensive substances of insects. Naturwissenschaften 64, 41–41. https://doi.org/10.1007/BF00439898

Evans, W.G., 1988. Chemically mediated habitat recognition in shore insects (Coleoptera: Carabidae; Hemiptera: Saldidae). J Chem Ecol 14, 1441–1454. https://doi.org/10.1007/BF01020147

Fockink, D.H., Mise, K.M., Zarbin, P.H.G., 2013. Male-Produced Sex Pheromone of the Carrion Beetles, Oxelytrum discicolle and its Attraction to Food Sources. J Chem Ecol 39, 1056–1065. https://doi.org/10.1007/s10886-013-0329-5

Francke, W., Schröder, F., Philipp, P., Meyer, H., Sinnwell, V., Gries, G., 1996. Identification and synthesis of new bicyclic acetals from the mountain pine beetle, Dendroctonus ponderosae hopkins (Col.: Scol.). Bioorganic & Medicinal Chemistry 4, 363–374. https://doi.org/10.1016/0968-0896(96)00013-2

Frérot, B., Foster, S.P., 1991. Sex pheromone evidence for two distinct taxa withinGraphania mutans (Walker). J Chem Ecol 17, 2077–2093. https://doi.org/10.1007/BF00987993

Gries, G., Borden, J.H., Pierce, H.D., Johnston, B.D., Oehlschlager, A.C., 1992. 3,7,7-trimethyl-1,3,5-cycloheptatriene in volatiles of female mountain pine beetles,Dendroctonus ponderosae. Naturwissenschaften 79, 27–28. https://doi.org/10.1007/BF01132277

Huth, A., Dettner, K., 1990. Defense chemicals from abdominal glands of 13 rove beetle species of subtribe staphylinina (Coleoptera: Staphylinidae, Staphylininae). J Chem Ecol 16, 2691–2711. https://doi.org/10.1007/BF00988079

Kim, J., Lee, S.-G., Shin, S.-C., Kwon, Y.-D., Park, I.-K., 2009. Male-Produced Aggregation Pheromone Blend in Platypus koryoensis. J. Agric. Food Chem. 57, 1406–1412. https://doi.org/10.1021/jf8032717

Kohnle, U., Mussong, M., Dubbel, V., Francke, W., 1987. Acetophenone in the aggregation of the beech bark beetle, *Taphrorychus bicolor* (Col., Scolytidae) ^1^. J Applied Entomology 103, 249–252. https://doi.org/10.1111/j.1439-0418.1987.tb00983.x

Lacey, E.S., Moreira, J.A., Millar, J.G., Hanks, L.M., 2008. A Male-produced Aggregation Pheromone Blend Consisting of Alkanediols, Terpenoids, and an Aromatic Alcohol from the Cerambycid Beetle Megacyllene caryae. J Chem Ecol 34, 408–417. https://doi.org/10.1007/s10886-008-9425-3

Lanne, B.S., Bergström, G., Wassgren, A.-B., Törnbäck, B., 1987. Biogenetic pattern of straight chain marking compounds in male bumble bees. Comparative Biochemistry and Physiology Part B: Comparative Biochemistry 88, 631–636. https://doi.org/10.1016/0305-0491(87)90355-5

Löfqvist, J., Bergström, G., 1980. Volatile communication substances in Dufour’s gland of virgin females and old queens of the antFormica polyctena. J Chem Ecol 6, 309–320. https://doi.org/10.1007/BF01402910

Molander, M.A., Eriksson, B., Winde, I.B., Zou, Y., Millar, J.G., Larsson, M.C., 2019. The aggregation-sex pheromones of the cerambycid beetles Anaglyptus mysticus and Xylotrechus antilope ssp. antilope: new model species for insect conservation through pheromone-based monitoring. Chemoecology 29, 111–124. https://doi.org/10.1007/s00049-019-00281-5

Morgan, E.D., Jungnickel, H., Billen, J., Ito, F., Bergmann, J., Gobin, B., 2008. Contents of the exocrine glands of the ant subfamily Cerapachyinae. Biochemical Systematics and Ecology 36, 260–265. https://doi.org/10.1016/j.bse.2007.02.007

Morgan, E.D., Parry, K., Tyler, R.C., 1979. The chemical composition of the Dufour gland secretion of the ant *Myrmica scrabrinodis*. Insect Biochemistry 9, 117–121. https://doi.org/10.1016/0020-1790(79)90036-2

Pianaro, A., Menezes, C., Kerr, W.E., Singer, R.B., Patricio, E.F.L.R.A., Marsaioli, A.J., 2009. Stingless Bees: Chemical Differences and Potential Functions in Nannotrigona testaceicornis and Plebeia droryana Males and Workers. J Chem Ecol 35, 1117–1128. https://doi.org/10.1007/s10886-009-9679-4

Pureswaran, D.S., Borden, J.H., 2004. New repellent semiochemicals for three species of Dendroctonus (Coleoptera: Scolytidae). Chemoecology 14, 67–75. https://doi.org/10.1007/s00049-003-0260-2

Pureswaran, D.S., Gries, R., Borden, J.H., 2004. Antennal responses of four species of tree-killing bark beetles (Coleoptera: Scolytidae) to volatiles collected from beetles, and their host and nonhost conifers. Chemoecology 14, 59–66. https://doi.org/10.1007/s00049-003-0261-1

Rochat, D., Meillour, P.N.-L., Esteban-Duran, J.R., Malosse, C., Perthuis, B., Morin, J.-P., Descoins, C., 2000. [No title found]. Journal of Chemical Ecology 26, 155–187. https://doi.org/10.1023/A:1005497613214

Scheffrahn, R.H., Gaston, L.K., Sims, J.J., Rust, M.K., 1984. Defensive Ecology ofForelius Foetidusand Its Chemosystematic Relationship toF. (=Iridomyrmex) pruinosus(Hymenoptera: Formicidae: Dolichoderinae). Environmental Entomology 13, 1502–1506. https://doi.org/10.1093/ee/13.6.1502

Steidle, J.L.M., Dettner, K., 1995. The chemistry of the abdominal gland secretion of six species of the rove beetle genus Bledius. Biochemical Systematics and Ecology 23, 757–765. https://doi.org/10.1016/0305-1978(95)00066-6

Teal, P.E.A., Mitchell, E.R., Tumlinson, J.H., Heath, R.R., Sugie, H., 1985. Identification of volatile sex pheromone components released by the southern armyworm,Spodoptera eridania (Cramer). J Chem Ecol 11, 717–725. https://doi.org/10.1007/BF00988301

Tengö, J., Bergstrom, G., 1977. Cleptoparasitism and Odor Mimetism in Bees: Do Nomada Males Imitate the Odor of Andrena Females? Science 196, 1117–1119.

Thiéry, D., Bonnard, O., Riquier, L., De Revel, G., Monceau, K., 2018. An alarm pheromone in the venom gland of Vespa velutina: evidence revisited from the european invasive population. entomologia 38, 145–156. https://doi.org/10.1127/entomologia/2018/0719

Torto, B., Suazo, A., Alborn, H., Tumlinson, J.H., Teal, P.E.A., 2005. Response of the small hive beetle ( *Aethina tumida* ) to a blend of chemicals identified from honeybee ( *Apis mellifera* ) volatiles. Apidologie 36, 523–532. https://doi.org/10.1051/apido:2005038

Tschinkel, W.R., 1975. A comparative study of the chemical defensive system of tenebrionid beetles: Chemistry of the secretions. Journal of Insect Physiology 21, 753–783. https://doi.org/10.1016/0022-1910(75)90008-6

Vinson, S.B., Williams, H.J., Frankie, G.W., Wheeler, J.W., Blum, M.S., Coville, R.E., 1982. Mandibular glands of maleCentris adani, (Hymenoptera: Anthophoridae): Their morphology, chemical constituents, and function in scent marking and territorial behavior. J Chem Ecol 8, 319–327. https://doi.org/10.1007/BF00987780

Wheeler, J.W., Happ, G.M., Araujo, J., Pasteels, J.M., 1972. γ-Dodecalactone from rove beetles. Tetrahedron Letters 13, 4635–4638. https://doi.org/10.1016/S0040-4039(01)94385-0
